# Environmental changes associated with drying climate are expected to affect functional groups of pro- and microeukaryotes differently in temporary saline waters

**DOI:** 10.1101/2022.11.29.518313

**Authors:** Zsuzsanna Márton, Beáta Szabó, Csaba F. Vad, Károly Pálffy, Zsófia Horváth

## Abstract

Temporary ponds are among the most sensitive aquatic habitats to climate change. Their microbial communities have crucial roles in food webs and biogeochemical cycling, yet how their communities are assembled along environmental gradients is still understudied. This study aimed to reveal the environmental drivers of diversity (OTU-based richness, evenness, and phylogenetic diversity) and community composition from a network of saline temporary ponds, soda pans, in two consecutive spring seasons characterized by contrasting weather conditions. We used DNA-based molecular methods to investigate microbial community composition. We tested the effect of environmental variables on the diversity of prokaryotic (bacteria, cyanobacteria) and microeukaryotic functional groups (ciliates, heterotrophic flagellates and nanoflagellates, fungi, phytoplankton) within and across the years. Conductivity and the concentration of total suspended solids and phosphorus were the most important environmental variables affecting diversity patterns in all functional groups. Environmental conditions were harsher and they also had a stronger impact on community composition in the dry spring. Our results imply that these conditions, which are becoming more frequent with climate change, have a negative effect on microbial diversity in temporary saline ponds. This eventually might translate into community-level shifts across trophic groups with changing local conditions with implications for ecosystem functioning.

## Introduction

Temporary ponds are a globally widespread habitat type with diverse aquatic communities and a wide variety of ecological functions ^1,2^. Despite their small size, their importance in terms of biodiversity is well-known for multiple groups including amphibians ^3–5^, macrophytes ^6–8^, and macroinvertebrates ^9,10^, and they also serve as a refuge for several rare or endangered species ^1,4^. At the same time, less is known about their microbial biodiversity^11^. Microorganisms not only provide the foundations of local food webs, but their contribution to trophic relationships and biogeochemical processes are essential in temporary aquatic ecosystems ^12,13^. Even so, biodiversity studies on microorganisms in temporary ponds are still much rarer than those on macroorganisms ^14^. As microbial biodiversity is linked to key ecosystem functions, such as nutrient cycling, mineralization ^15,16^ and ecosystem services ^17,18^, identifying its drivers is of high importance to understand the functioning of temporary waters.

Environmental harshness is generally higher in temporary ponds than permanent ones due to periodic droughts and rapid seasonal changes in environmental conditions ^19–21^. As temporary aquatic habitats are usually very shallow, local species often need to withstand high UV radiation ^22^ and extreme daily fluctuations in temperature ^23^. Saline temporary waters represent even more extreme habitats, where the overall high salinity levels and their seasonal fluctuation represent additional environmental stress ^24^. Microorganisms are known to be able to sustain large local population sizes even in such extreme conditions due to their high physiological and evolutionary adaptation potential linked to their small size and short generation times ^11,25^. Even so, saline inland waters have so far received little attention when it comes to the diversity of microbial communities ^26^ compared to estuaries ^27–29^ and man-made salterns ^30–32^. Consequently, less is known about temporary saline waters and existing studies are generally limited to specific groups like diatoms ^33–35^ or bacteria ^36,37^, based on small spatial scales and a limited number of habitats ^26^. Although salinity has been a focal point of studies ^38–40^, we still have little information about the phylogenetic diversity of microorganisms along salinity gradients in temporary ponds and how community composition or the relative share of multiple functional groups change along these gradients. Regional-scale studies could deliver important insights into these patterns and processes, but they are still largely missing ^26^.

Shortening hydroperiods represent a global trend across multiple regions, with naturally permanent ponds becoming temporary ^43^ and with temporary ponds shrinking gradually and eventually disappearing ^4,41,42^. Several regions with naturally abundant temporary ponds and wetlands have already suffered significant habitat losses ^42,46,47^. The shortening hydroperiod of temporary ponds under climate change is expected to result in loss of biodiversity ^44,45,48^, especially when coupled with other stressors, such as an increased rate of salinization ^88^. Increasing salinity has been shown to alter the structure of microbial communities in ponds ^26,40^, which has a cascading effect on ecosystem functions, e.g., the balance between greenhouse gas emission and carbon sequestration ^49–51^. At the same time, studies on the responses of microbial communities to elevated salinities were mostly done in the context of salinization of freshwaters ^52^. Therefore, our knowledge of how microbial communities of naturally saline habitats will respond to such changes is still limited.

Soda lakes are a special type of saline lakes, with carbonate (CO32-), bicarbonate (HCO3-) and sodium (Na^2+^) as the dominant ions and a stable alkaline pH ^12,53^. They can be found worldwide but are less frequent than other inland saline aquatic habitats ^74^. In Europe, they mainly occur as temporary soda pans on the Pannonian Plain (Austria, Hungary, and Serbia) ^23^. These unique and regionally restricted ecosystems serve as important refuges for a number of rare or endangered species ^71,72^ and as important feeding sites for waterbirds ^73, 98^, but their numbers are decreasing ^47^. The long-term future of these vulnerable and geographically restricted inland saline ecosystems is threatened by the intensifying effects of climate change in the form of decreasing precipitation and increasing temperature in the region (Figure 1).

**Figure 1.**
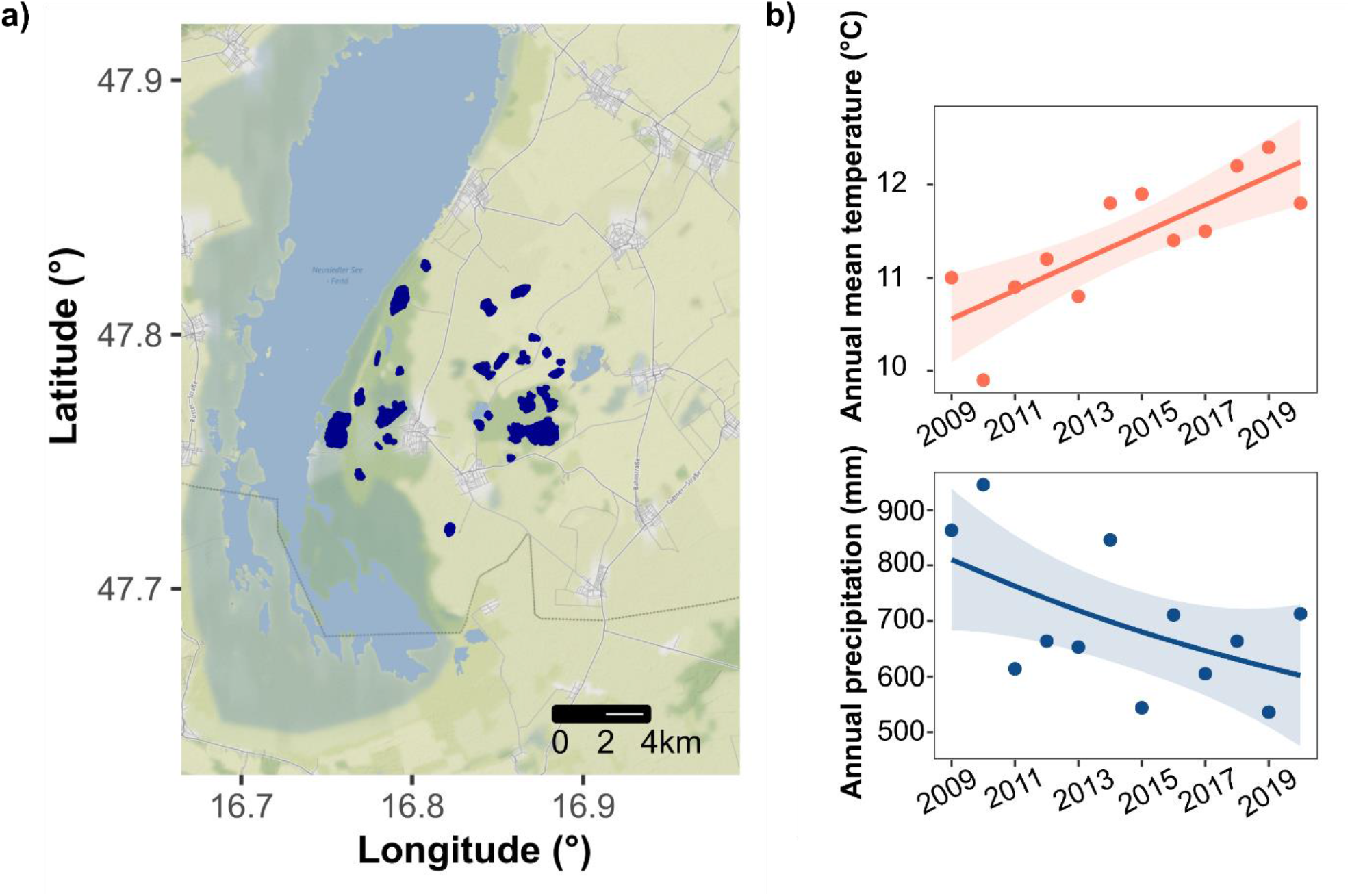
a) Location of the sampled soda pans (dark blue) in the National Park Neusiedlersee-Seewinkel in eastern Austria (Map was created with R v. 4.2.2 ^96^) b) Temporal trends in annual mean temperature and precipitation since 2009 based on meteorological data from the Seewinkel region (Station Eisenstadt; trendlines and confidence intervals are based on Generalized Additive Models, see also in Supplementary material Text S1)

Here, we present a study based on a network of natural saline temporary ponds to investigate diversity as well as community composition of pro-and microeukaryotes along local environmental gradients in two consecutive spring seasons with contrasting precipitation. Our study area is a network of temporary soda pans in eastern Austria, with salinities that can range from sub-(0.5-3 g/L) to hyposaline (3-20 g/L) values ^53^. Due to the close proximity of the ponds, they are under the same climatic and geographical conditions, making them a suitable regional case study for the environmental drivers of microbial communities. Specifically, our first aim is to explore the main environmental drivers of taxonomic and phylogenetic richness and composition of microorganism communities (functional groups of pro-and microeukaryotes). Second, we test whether the identity and strength of the main drivers change with contrasting climatic conditions (by comparing the dry spring of 2017 and the wetter 2018), which can have important implications for the future of these vulnerable ecosystems.

## Results

The two sampling years differed considerably based on the local environmental variables. Water level was significantly lower in 2017 than in 2018, which coincided with significantly higher values of conductivity, TN, TP, TSS, and pH (Figure 2).

**Figure 2.**
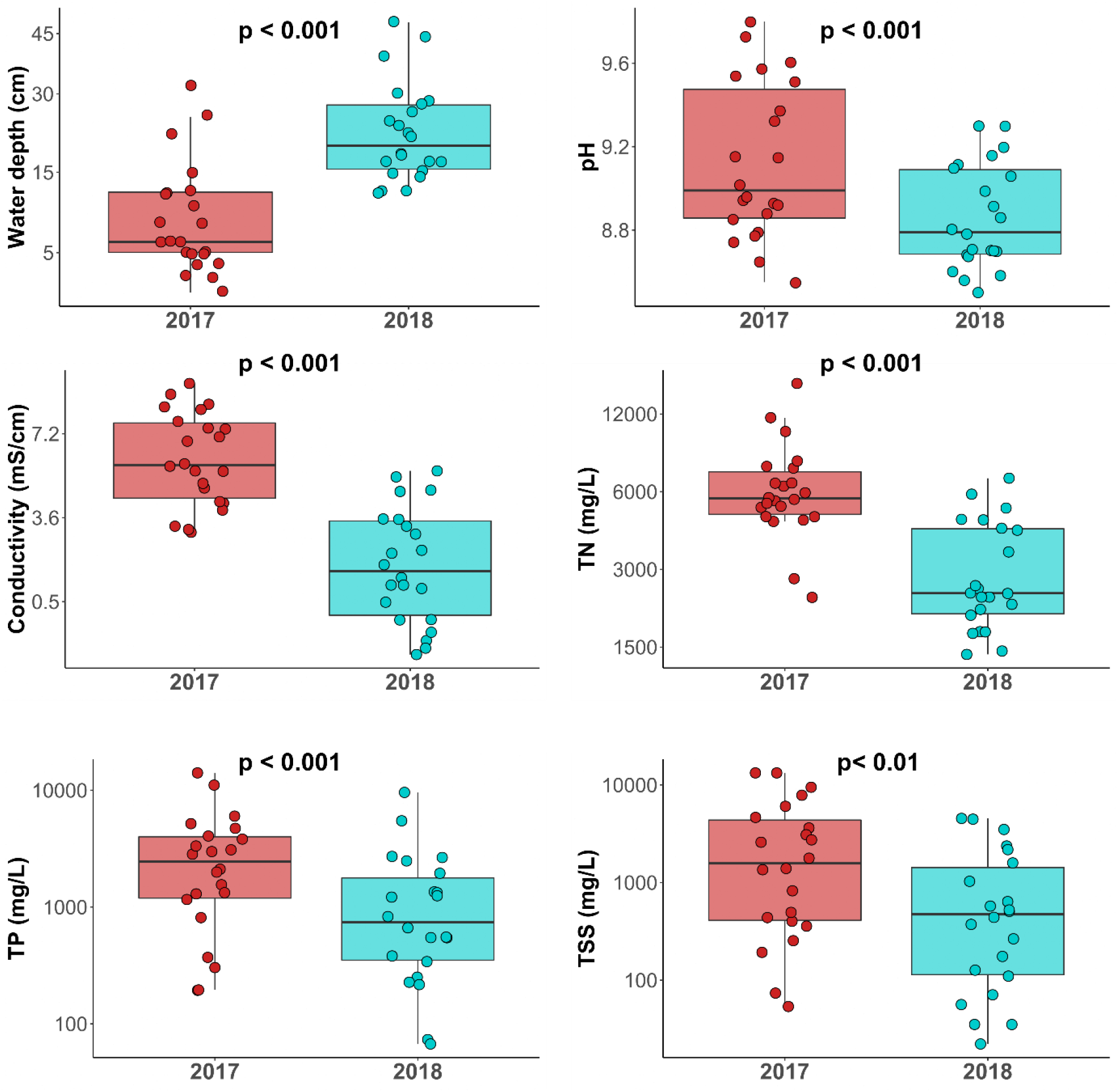
Boxplots illustrating differences in the environmental variables between the two sampling years. *P* values indicate significant differences between years based on paired t-tests.

Based on the comparison of environmental variables with reference dates (according to a pairwise PERMANOVA and NMDS; Table 1, Figure 3), the wetter spring of 2018 was comparable to spring conditions from a decade ago (2010), while it differed significantly both from the dry spring (2017) and the reference summer conditions (2009). On the other hand, dry spring conditions (2017) did not differ statistically from summer conditions (2009).

**Table 1.** Results of the pairwise comparisons of the reference data and our data via PERMANOVA between years (2017 spring, 2018 spring data and 2009 summer, 2010 spring) reference data (p values are adjusted for multiple comparisons based on Bonferroni method, p < 0.05, bold: significant)

| Pairs |  | Df | SS | F | R<sup>2</sup> | p adjusted |
| --- | --- | --- | --- | --- | --- | --- |
| 2009 summer | 2010 spring | 1 | 3.036 | 0.998 | 0.0322 | 1.000 |
| 2009 summer | 2017 spring | 1 | 4.964 | 1.853 | 0.0582 | 1.000 |
| **2009 summer** | **2018 spring** | **1** | **24.136** | **9.776** | **0.246** | **0.018** |
| 2010 spring | 2017 spring | 1 | 9.718 | 4.185 | 0.122 | 0.153 |
| 2010 spring | 2018 spring | 1 | 10.750 | 5.089 | 0.145 | 0.051 |
| **2017 spring** | **2018 spring** | **1** | **30.977** | **17.697** | **0.371** | **0.003** |

**Figure 3.**
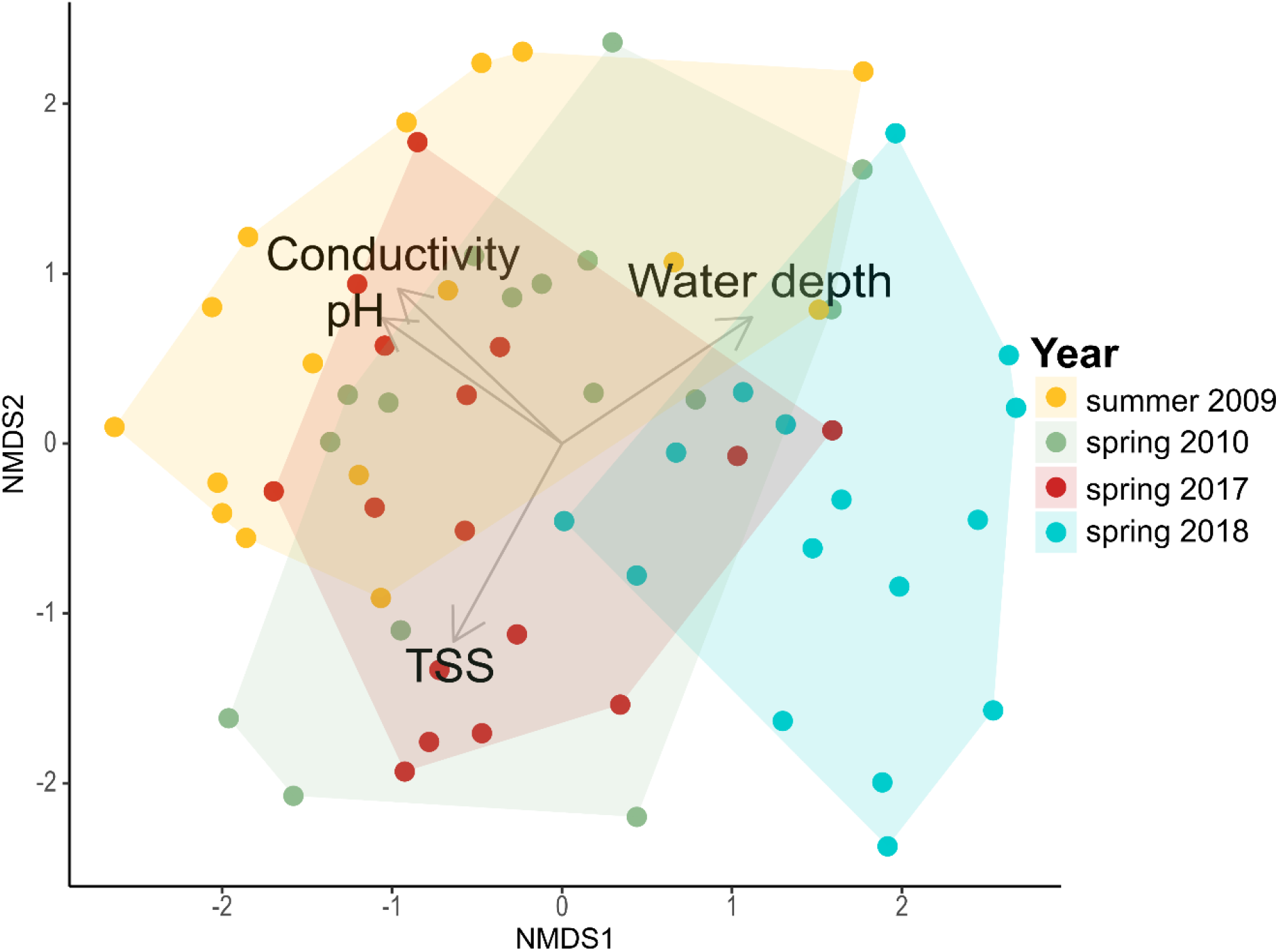
NMDS plot illustrating the separation of samples (each soda pan is represented by one filled circle per year) based on the local environmental variables in the Seewinkel regional data (2017: dry, 2018: wetter) and the earlier reference data (2009: summer, 2010: spring)

Richness of bacteria and cyanobacteria was higher in 2017 than in 2018, which was also reflected by the significantly higher PD (phylogenetic diversity) in these two groups in 2017 (Figure 4). Bacteria also showed significantly higher evenness values in 2017 along with ciliates. Phytoplankton showed different patterns, with higher evenness and PD, but lower richness in 2017 (Figure 4). The rest of the indices - ciliate richness and PD and all diversity indices of fungi and HF-HNF (heterotrophic flagellates and nanoflagellates - showed no difference between the two years.

**Figure 4.**
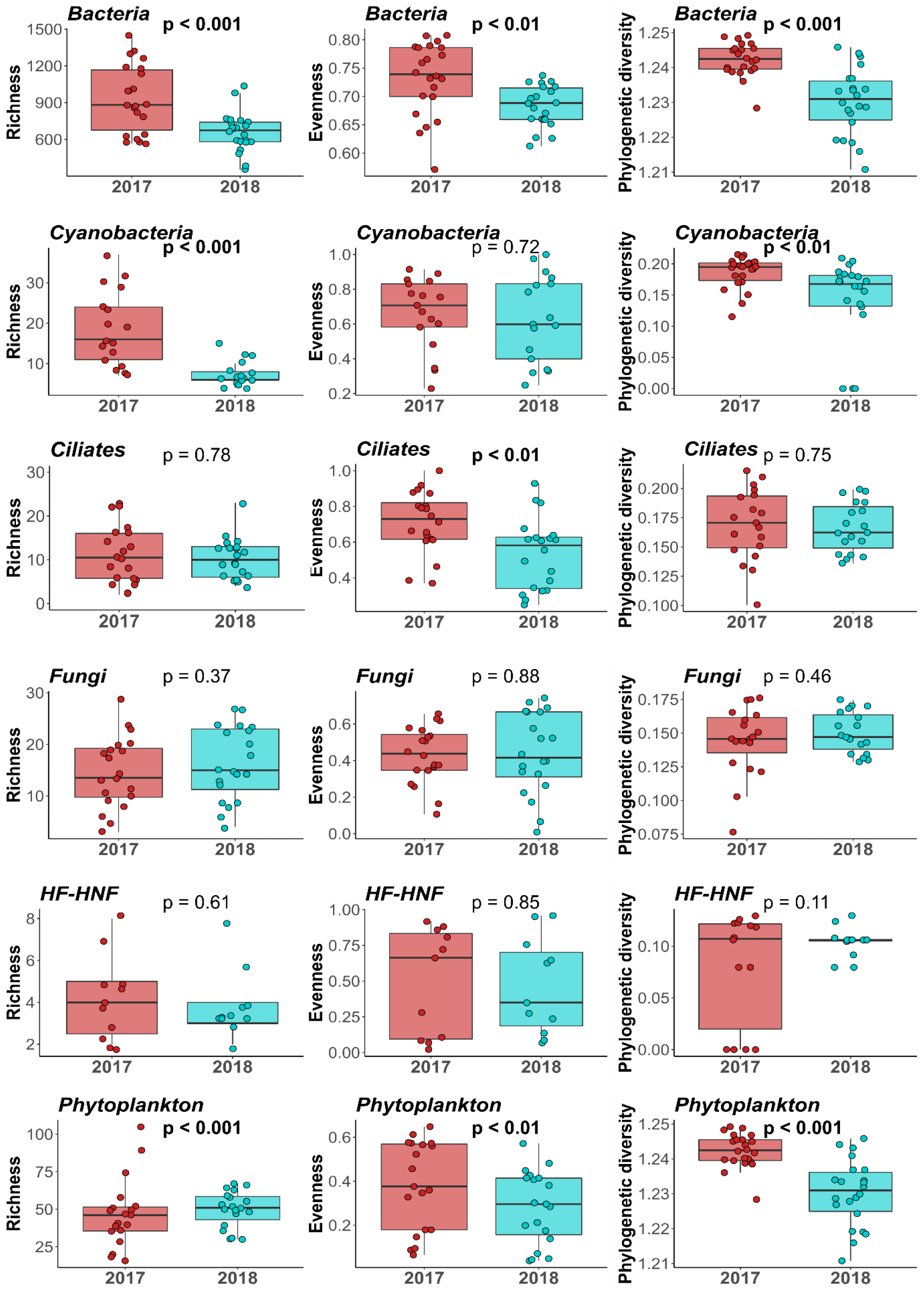
Richness, evenness and phylogenetic diversity of the microeukaryotic and prokaryotic groups in the two sampling years. Significance values are based on paired t-tests (bold: significant)

Prokaryotic phylum composition was significantly influenced by water depth and TP in both years, with conductivity being additionally significant in 2018 (Figure 5a, Table S4). Among the prokaryotic phyla, photosynthetic cyanobacteria were more abundant in more saline and deeper pans which was more explicit in 2018 (Figure 5). For microeukaryotic phyla, conductivity was a significant predictor in both years according to the CCAs (Figure 5b), together with either TP (2017) or pH (2018) (Table S5). A separation of taxa along the conductivity gradient was conspicuous in both years, with mixotrophic (Dinoflagellata and/or Cryptophyta) and heterotrophic groups (Centroheliozoa, Alveolata_unclassified, Ciliophora) being more abundant in less saline waters (i.e., of lower conductivity), while the relative abundances of Haptophyta and Discoba increased with conductivity (Figure 5). Microeukaryotic functional groups were significantly influenced only by conductivity in 2017, while in 2018, no environmental variable was significant (Figure S3). A separation of the functional groups along conductivity can be seen in both years, with the relative abundance of ciliates decreased with conductivity, while phytoplankton, fungi, and particularly heterotrophic flagellates and nanoflagellates (HF-HNF) were more abundant in more saline waters.

**Figure 5.**
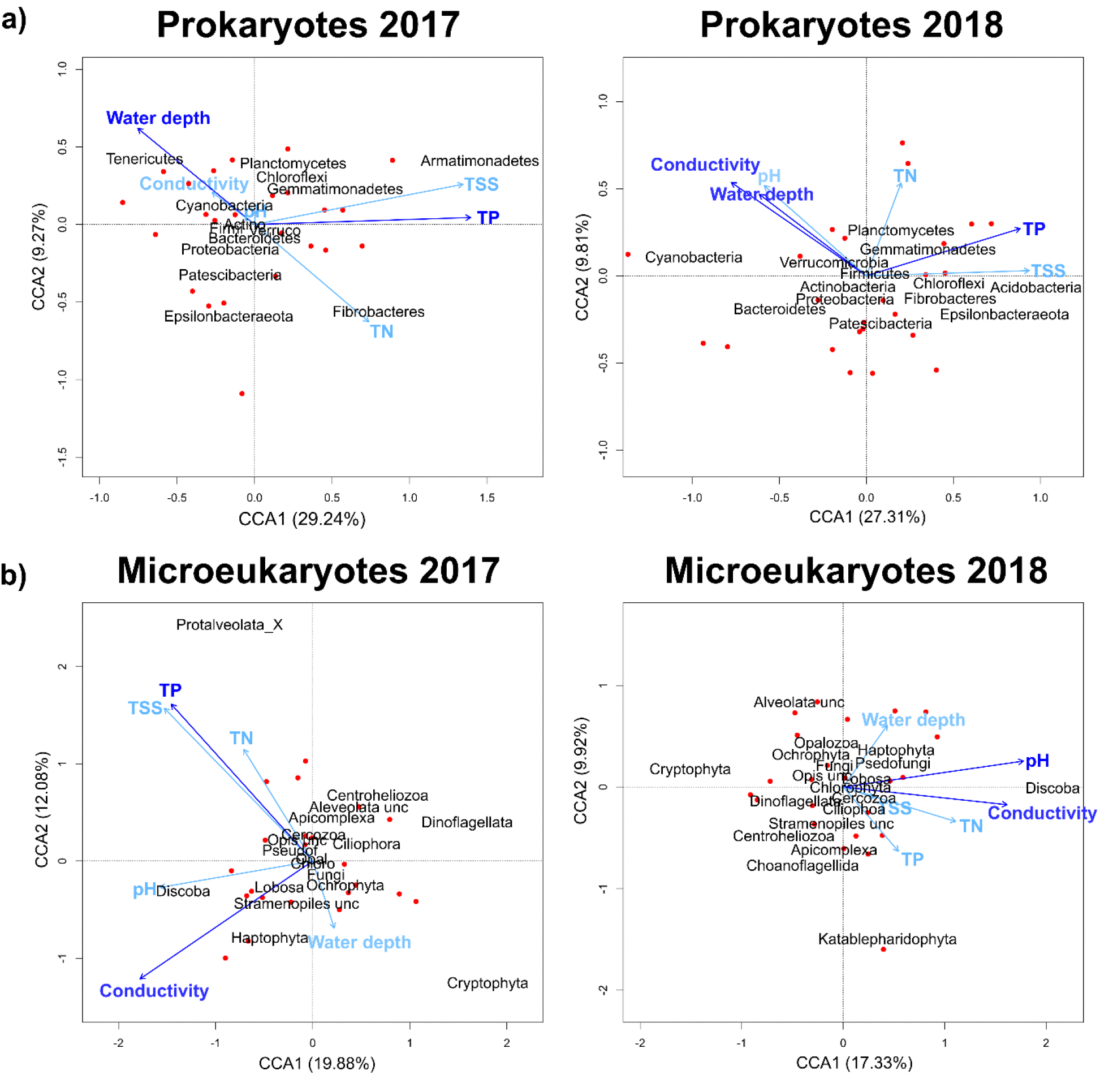
Canonical correspondence analysis (CCA) biplots based on the phylum composition of prokaryotes (a) and microeukaryotes (b) in the two years (red points: habitat scores, dark blue arrows: significant environmental variables, light blue arrows: all the other environmental variables). Abbreviation of the phyla names: unc = unclassified, Opis unc = Opisthokonta unclassified, Firmi= Firmicutes, Verruco = Verrucomicrobia, Actino = Actinobacteria, Chloro = Chlorphyta, Psedof = Pseudofungi, Ochro = Ochrophyta, Opal = Opalozoa

Water depth, conductivity and TP were similarly important predictors of OTU composition in the separate CCA analyses of the major functional groups (Table 2). Conductivity was selected as a significant predictor in almost all models. There was a general trend that the environmental variables explained a higher variation in OTU composition in 2017 (explained variation ranging between 0.12 - 0.25) than in 2018 (0 - 0.15).

**Table 2.** Significant environmental variables (p < 0.05) of the OTU composition of prokaryotic and microeukaryotic functional groups based on separate CCAs (presented in the supplementary material, Figure S1-2)

|  |  | Water<br>depth | pH | Conductivity | TN | TP | TSS | Explained<br>variation |
| --- | --- | --- | --- | --- | --- | --- | --- | --- |
| 2017 |  |  |  |  |  |  |  |  |
| Prokaryotes | Bacteria | ● |  | ● |  | ● |  | 0.16 |
|  | Cyanobacteria | ● |  | ● |  | ● |  | 0.15 |
| Microeukaryotes | Ciliates |  |  |  |  | ● | ● | 0.16 |
|  | Fungi | ● |  | ● |  | ● |  | 0.14 |
|  | HF-HNF |  |  | ● |  |  |  | 0.12 |
|  | Phytoplankton | ● |  | ● | ● | ● |  | 0.25 |
| 2018 |  |  |  |  |  |  |  |  |
| Prokaryotes | Bacteria | ● |  | ● |  | ● |  | 0.15 |
|  | Cyanobacteria |  |  |  |  |  |  | NA |
| Microeukaryotes | Ciliates |  |  | ● |  |  |  | 0.06 |
|  | Fungi |  |  | ● |  |  |  | 0.04 |
|  | HF-HNF |  |  | ● |  |  |  | 0.07 |
|  | Phytoplankton | ● |  | ● |  |  |  | 0.12 |

Local species richness of the six major groups (bacteria, cyanobacteria, ciliates, fungi, HF-HNF, phytoplankton) generally responded linearly to environmental gradients. In case of evenness, several non-linear relationships were observed (e.g., GAM predictions provided a better fit over linear models for five out of the six models for evenness in 2018; Figure 6). We also found U-shaped patterns in a few cases, including bacteria (PD with TSS) (Figure S4), ciliates (richness with TSS, evenness with water depth) (Figure S6), fungi (richness with pH) (Figure S7), HF-HNF (evenness with TP) (Figure S8), and phytoplankton (PD with conductivity and pH) (Figure S9).

**Figure 6.**
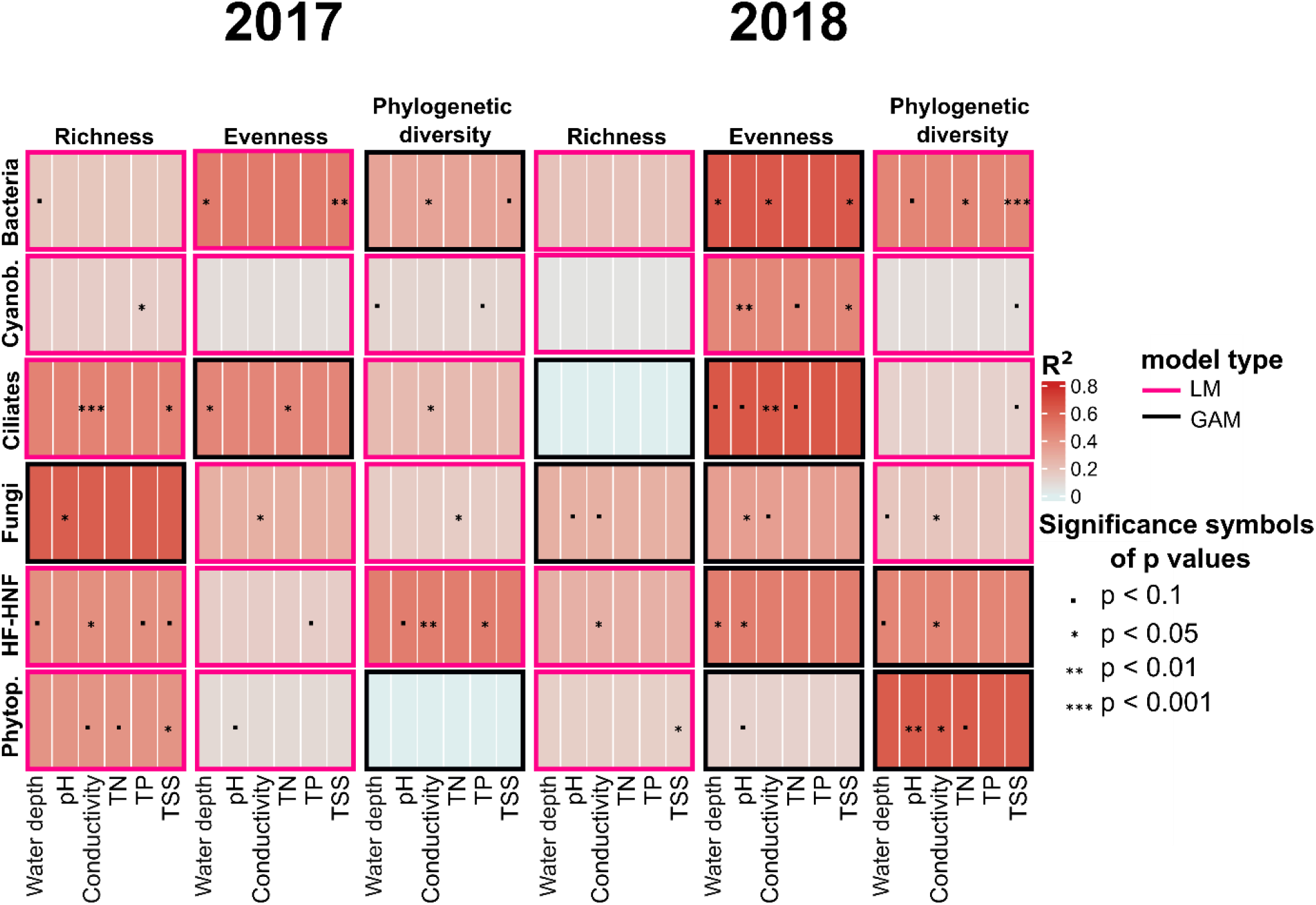
Heatmap of the best-fit models with the significant environmental variables for the six major taxonomic groups (in rows) and their local diversity indices (richness, evenness, phylogenetic diversity) in the two sampling years (in columns). One model is represented by one box, where the shading indicates the overall R^2^ of each multiple regression model (either linear or GAM-based, indicated by the colour of the frame of each box), while the significance symbols belong to individual predictors in the final model. Abbreviation of the group names: Cyanob. = Cyanobacteria, Phytop. = Phytoplankton, HF-HNF = Heterotrophic flagellates and nanoflagellates

Prokaryote richness (bacteria and especially cyanobacteria) was in general poorly explained by the environment (R^2^ of the LMs of bacteria: 0.175 in 2017 and 0.189 in 2018, R^2^ of LM of cyanobacteria: 0.141 in 2017 and 0.05 in 2018), and no significant predictor was identified for either group in 2018. In general, environment explained a higher variation in richness in 2017 than in 2018 in all microeukaryotic groups and in cyanobacteria (based on R^2^ values of the models, Figure 6).

Conductivity and TSS were the most frequent significant predictors for richness and PD across the taxonomic groups (Figure 6). Among all the functional groups, HF-HNF was the only one with a positive relationship with conductivity, which was significant both for richness and PD in both years (Figure S8). Bacteria and cyanobacteria generally showed negative trends with conductivity (or negative after a threshold) within the two years, which changed to an overall positive trend when the two years were analysed together (Figure S4-5). The most frequent significant predictor of evenness was pH across the different groups, while water depth and conductivity were also found to be significant for some of the functional groups (Figure 6).

We found that the relationship between changes in local richness and environmental variables were significant in three out of the six functional groups (Table S6). Local changes in bacteria and ciliate richness corresponded significantly to changes in conductivity, while for fungi, the main driver was the change in pH between the two years. These responses were all negative, indicating a drop in richness with increasing conductivity and pH (Figure 7).

**Figure 7.**
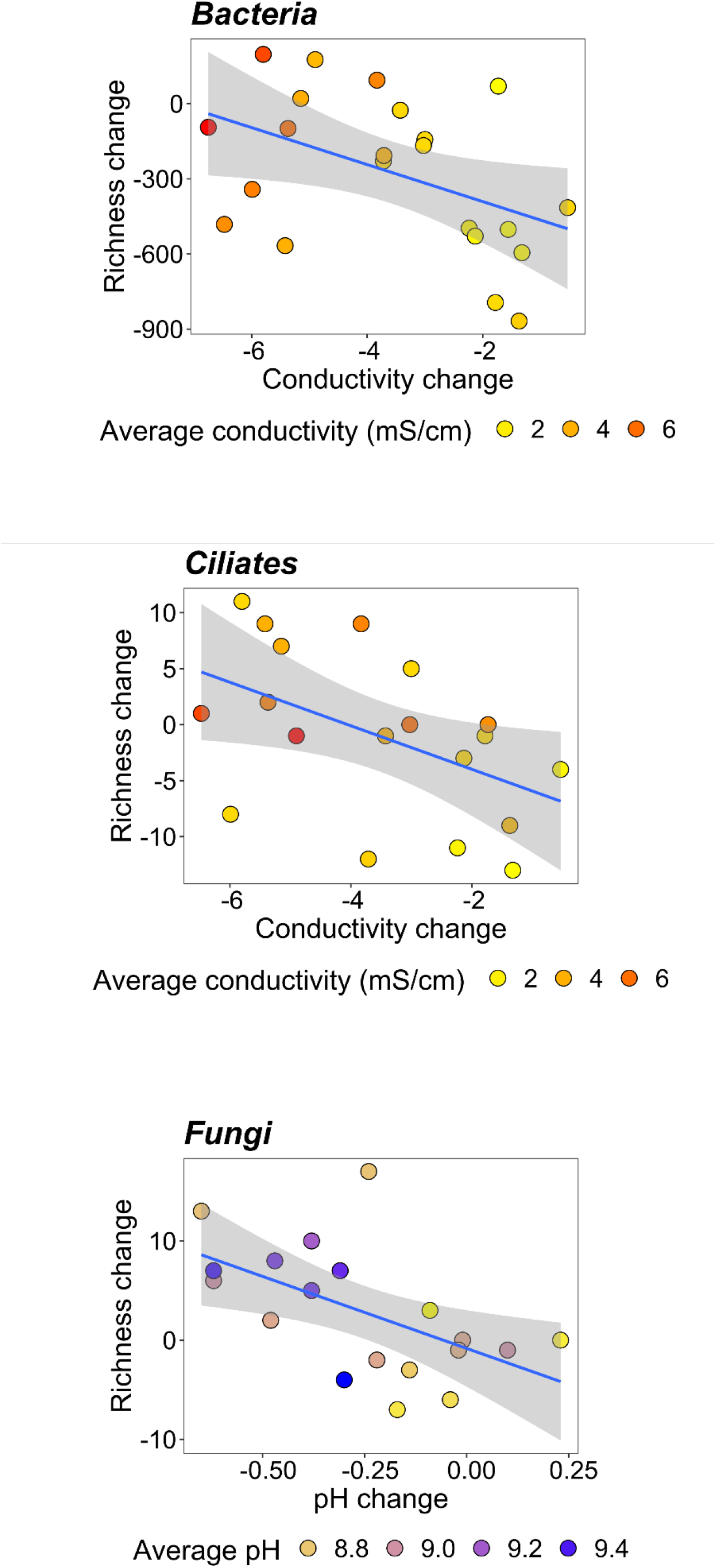
The relationship between richness changes and significant environmental variable changes (p < 0.01)

## Discussion

We analysed diversity patterns of microbial communities along environmental gradients in two consecutive spring seasons with contrasting weather and hydrological conditions. While environmental conditions in the wetter spring (2018) were comparable to the reference spring data from a decade ago, spring conditions in the dry year (2017) were more similar to the reference summer data, indicating a seasonal shift in the aquatic environment. A general trend was found that environmental variables had stronger effects on the community composition in the dry spring. Conductivity, TSS, and TP were the most important environmental variables of the diversity and community patterns across six major functional groups of microbes. At the same time, the response of prokaryotes (bacteria and cyanobacteria) to the environmental gradients generally differed from the microeukaryotic functional groups. With the ongoing climate change, drier weather conditions and stronger environmental stress gradients can be expected in the region, much like in our 2017 dataset. Therefore, climate change might drive shifts in community composition of microbial communities in temporary saline ponds, where microeukaryotic plankton may be more adversely affected by the increasing environmental stress.

According to our results, conductivity was overall the most important environmental variable shaping microbial diversity patterns. Even though the salinity gradient of the studied habitats was relatively short (ranging from 0.6 to 11 mS/cm conductivity, which equals to 0.4-8.8 g/L salinity based on a previously published conversion factor ^53^), studies indicate that the major changes in saline lake communities are rather expected to happen at these lower salinity levels, with major shift between 3-10 g/L ^54–56^. A former study encompassing all soda pans in the Pannonian ecoregion found a threshold of 3.9 g/L, above which a more pronounced drop in zooplankton species richness was observed ^39^. Our data was in agreement with this for most microbial groups, as above 3.2 mS/cm conductivity (2.6 g/L salinity), we observed negative trends in richness in five of the six groups (except for HF-HNF which showed a strikingly different pattern, with a clear increase in richness with conductivity). As most conductivity values were higher than 3 mS/cm in 2017, this can also explain the relatively lower biodiversity values for most groups in 2017 compared to 2018.

The high inorganic turbidity of the pans makes traditional microscopic identification of microeukaryotes challenging in these systems. This might explain the general lack of community-level studies on microeukaryotes from these habitats. Our study is the first regional survey based on molecular data of the 18S rRNA gene, which resulted in the first high resolution dataset for multiple microeukaryote functional groups in soda pans. In phytoplankton, the dominance of the pico-sized fraction is also a typical phenomenon in these habitats, where traditional microscopic techniques offer limited taxonomic resolution ^57–59^. Accordingly, a previous review based on microscopic data from soda pans found species numbers rarely exceeding ten ^60^. Compared to this, we found an average eukaryotic phytoplankton OTU richness of 47 in 2017 and 49 in 2018, with the lowest value of 16 (Runde Lacke in 2017) and the highest value of 105 (Obere Höllacke in 2017). These findings show that these habitats harbour a much higher phytoplankton diversity than previously assumed.

Phytoplankton was the only group with a U-shaped pattern in phylogenetic diversity along the salinity gradient in 2018 (Figure S9). Intermediate salinity (2.9 g/L) had the lowest phylogenetic diversity, beyond which phylogenetic diversity increased again with salinity. Similar U-shaped species richness patterns of phytoplankton along salinity gradients are recorded from transitional gradients from freshwater through brackish to marine habitats ^61,62^. In soda pans, however, the increase in phylogenetic diversity was not coupled with a parallel increase in OTU richness, which suggests that this pattern rather emerged due to the disappearance of closely related taxa rather than the emergence of salt tolerant OTUs.

HF-HNF richness and phylogenetic diversity clearly showed an increasing trend with salinity. Heterotrophic nanoflagellates are the main protozoan grazers of bacterio- and picophytoplankton ^63^. Previous studies from other saline systems showed that heterotrophic flagellate diversity is mainly driven by salinity and temperature ^64,65,66^ and follows the “rule of critical salinity”, which means species richness has its minimum in brackish waters (~5-8 g/L of salinity) ^67^. In our study, HF-HNF richness and phylogenetic diversity showed an exponential increase with increasing salinity, which is a pattern previously not reported from other saline systems. Further studies will be necessary to explore the relative effect of salinity and the possible indirect effects through changes in biotic interactions underlying this pattern.

Even though the environmental variables did not constrain all taxa the same way in the microbial communities, our results indicated that environment explained more community variation in the dry spring (2017), which is in agreement with species sorting becoming stronger with harsher environmental conditions ^68–70^. The response of biodiversity indices to changes in environmental conditions between the two years indicated that elevated levels of salinity and other parameters can be associated with lower levels of biodiversity for most of the microbial functional groups. The limitation of our study is that we cannot compare our recent data to earlier community datasets as this is the first regional survey on the microbial communities of these ponds based on amplicon sequencing. Although we hold no long-term data, our results indicate that climate change can be expected to have a serious negative impact on the biodiversity of these aquatic habitats. We promote that longer term monitoring studies would be necessary to follow the long-term changes in these unique aquatic habitats, to explore the direct links between the changing climatic conditions and their biota.

## Conclusions

Our results demonstrated that soda pans host diverse microbial communities, which were so far largely unexplored due to the lack of regional studies and the limited application of molecular methods. The ongoing environmental changes will most likely lead to biodiversity loss in multiple microbial groups according to our results, which might induce critical ecosystem-level consequences. Therefore, longer term biodiversity studies and targeted research for better understanding the functional role of the most sensitive microbial groups would be crucial, not only in this type of ecosystem, but also in other temporary saline pond networks.

## Methods

### Sampling and sample processing

Water samples were collected from 26 soda pans from the region of National Park Neusiedlersee-Seewinkel, Austria in the same spring period of 2017 (3-6 of April) and 2018 (2-4 of April) (Figure 1, Table S1). These two years had very contrasting weather conditions and hydroregimes. For the spring filling of soda pans, precipitation during the winter months is crucial ^24^. During the four months prior to our 2017 survey, the amount of average monthly precipitation (26 mm) was almost half of what was measured in 2018 (47 mm; Supplementary material, Table S2). Additionally, an increasing trend in annual temperature can be observed in the region (Figure 1), which likely contributes to higher evaporation rates. Altogether, there was an overall difference in the hydrological conditions, with one of the soda pans (Kirchsee) even being completely dry in April 2017.

Water depth and Secchi depth were measured in the open water of each soda pan. A composite water sample was collected from at least 20 different points of each soda pan using a one-liter beaker (total volume: 20 L). It was filtered through a plankton net (mesh size of 100 µm) to remove larger zooplankton and filamentous algae, to prevent sequencing bias in community composition assessments. 1 L of this filtered composite water sample was delivered to the laboratory in a glass bottle in a cool box for measuring total suspended solids (TSS) and processing molecular samples. We collected a similar but unfiltered composite water sample to measure conductivity and pH with a multimeter and to take samples for determining total nitrogen (TN) and total phosphorus (TP) concentration, which were subsequently frozen until further processing.

To measure the amount of TSS, water was filtered through a pre-weighted GF/F filter until it was clogged. The filters were dried overnight at 60 °C and they were weighed again to calculate TSS concentration. TN and TP were measured from the unfiltered water samples ^75, 89^. For molecular biology analysis, 1-50 mL of the composite water samples (depending on turbidity) were filtered through a nitrocellulose membrane filter (Ø 47 mm, 0.22 μm pore size). Filters were stored at −20 °C until DNA isolation.

### DNA extraction and sequencing

DNA was extracted from the filters by using PowerSoil DNA Isolation Kit (MO BIO Laboratories Inc., Carlsbad, CA, USA) according to the manufacturer’s instructions. Extracted DNA was stored at −20 °C until shipping to LGC Genomics (Berlin, Germany) for amplicon sequencing. For the determination of prokaryotic and microeukaryotic community composition, the V4 region of the 16S rRNA gene and the V7 region of the 18S rRNA gene were amplified, using prokaryotic primers EMBf 515F (GTGYCAGCMGCCGCGGTAA ^76^) – EMBr 806R (GGACTACNVGGGTWTCTAAT ^77^) and eukaryotic primers UnivF-1183mod (AATTTGACTCAACRCGGG) – UnivR-1443mod (GRGCATCACAGACCTG ^78^). The V7 region was chosen as a good compromise between fragment length (260-360 bp) and taxonomic resolution ^78, 97^. All polymerase chain reactions and amplicon sequencing (Illumina MiSeq platform) were carried out by LGC Genomics (Berlin, Germany) ^73^ (Table S3).

### Amplicon sequence analysis, taxonomic assignment, and compilation of community matrices

Bioinformatic analysis of the sequences were carried out with mothur v1.43.0 ^79^ using the MiSeq SOP http://www.mothur.org/wiki/MiSeq_SOP ^80^, downloaded on 12th November 2020, with the modification of adjusting the deltaq parameter to 10 in the “make.contigs” command. Primers were removed from both ends of the sequences and singletons were excluded from the dataset ^81^. Read alignments and taxonomic assignments were carried out using the ARB-SILVA SSU Ref NR 138 reference database with a minimum bootstrap confidence score of 80 ^82^. Reads assigned to non-target specific lineages (e.g., Archaea, Chloroplast, Mitochondria, unknown) were removed from the dataset. Operational taxonomic units (OTUs) were picked at 99% similarity threshold levels. For the 16S rRNA OTUs, TaxAss software ^83^ was used with default parameters based on the FreshTrain (15 June 2020 release) and ARB-SILVA SSU Ref NR 138 databases, the taxonomic assignment of 18S rRNA OTUs was performed with the PR2 v4.12.0 reference database ^84^. OTUs belonging to taxa Streptophyta, Metazoa, Ascomycota and Basidiomycota were excluded from the microeukaryotic dataset. Both the 16S rRNA gene and 18S rRNA gene datasets have been subsampled (rarified) to the lowest read count (8620 reads per sample for the 16S rRNA dataset and 2432 reads per sample for the 18S rRNA dataset). Hereinafter, the resulting dataset is referred to as relative abundance data.

The rarefied prokaryotic and microeukaryotic data were further split into groups, hereinafter referred to as functional groups, based on taxonomy (according to higher categories) and function. Delineation of groups based on taxonomy alone does not always reflect functional differences, while classification based on function may result in polyphyletic groups, therefore we opted for combining both types of information. This resulted in two groups of prokaryotes (16S rRNA data): heterotrophic bacteria and autotrophic cyanobacteria; as well as four groups of microeukaryotes (18S rRNA data): ciliates, fungi, phytoplankton, and heterotrophic flagellates and nanoflagellates (HF-HNF). The latter four groups were defined based on taxonomy and trophic position (i.e., autotrophic and heterotrophic groups). The functional groups were created with the subset_taxa function in the “phyloseq” package ^85^. Data are openly available in the NCBI SRA database as part of the BioProject accession PRJNA748202 ^73^.

### Statistical analysis

A set of environmental variables were available from a decade earlier, which included spring data from 2010 and summer data from 2009 ^23^ for 16 pans which we also sampled during our study (in 2017 and 2018). We used these data from 2009-2010 as a reference from a period still characterized by lower annual mean temperature and higher amount of mean annual precipitation in the region (Figure 1). To assess the overall differences in environmental conditions between the four timepoints, we first applied non-metric multidimensional scaling (NMDS). To normalize the distribution of all environmental variables, log transformation was used for conductivity, TP, TN, and TSS, and square root transformation was applied for water depth data prior to all analyses. We standardized all data to unit variance and calculated a Euclidean distance matrix of all normalized environmental data with the “vegan” package ^90^. We then tested the differences between the four years, for which we applied pairwise comparisons with a permutational multivariate analysis of variance (PERMANOVA) using the “pairwiseAdonis” R package by adjusting for multiple comparisons ^92^, using the same distance matrix.

To test for differences between the two years of our present study (2017 and 2018), we applied paired t-tests for each environmental variable using the package “ggpubr” ^91^. For the paired t-tests, data from only those habitats were included which were sampled in both years.

Richness (number of OTUs observed), Pielou’s evenness (referred as evenness later on)were used to measure the structure of the communities and phylogenetic diversity (PD) was used as a measure of functional diversity in the two sampling years. Richness and evenness were calculated using the “phyloseq” package ^85^. Phylogenetic trees were created with the FastTree software (Qiime2 plugin) ^86^. Phylogenetic diversity was quantified through Rao’s quadratic entropy from the R package “picante” ^93^. The three measures of diversity were compared in the two sampling years with paired t-tests on a subset of samples that were sampled in both sampling years.

Canonical correspondence analysis (CCA) was performed to examine the relationship between environmental variables and community composition. For prokaryotes and microeukaryotes, CCAs were created based on phyla, while relative abundances of OTUs were used for the six major functional groups (bacteria, ciliates, cyanobacteria, fungi, HF-HNF, phytoplankton) separately (using the package “vegan” ^90^). Additional CCAs were created based on the relative abundance of the microeukaryotic functional groups (not carried out for prokaryotes, represented only by two groups). Relative abundance data of prokaryotes and microeukaryotes were log(x+1) transformed prior to analysis. To reduce the complexity of the datasets, only those prokaryotic and microeukaryotic phyla were included in the analysis that had more than three occurrences and reached at least five percent relative abundance each year. For the six functional groups, OTU tables were used for the analysis (rare OTUs with less than four occurrences were excluded). Significant environmental variables were identified based on both backward and forward stepwise model selection using permutation tests (n = 199). In order to determine what proportion of total variation was explained by significant variables, CCAs for each group and year were rerun as partial CCAs, in which only the significant ones were kept as constraining variables, while all the others were partialled out as conditioning variables. The ratio of constrained and total inertia yielded the proportion of variation explained.

To analyze which environmental variables had the strongest effect on local diversity (richness, evenness, and PD), we compared the fit of multiple linear regression models (with the “MASS” package of R ^94^) and generalized additive models with smooth terms (using the “mgcv” package of R ^95^). Model selection was based on the Akaike’s Information Criteria (AIC). Separate models were fitted for the local diversity indices of the six taxonomic groups and three time periods: data of the two sampling years (2017, 2018) separately or pooled, resulting in a total of 36 multiple regression models. We created a summary table of the best fitted ML or GAM models for the six functional groups in the two years with the significant environmental variables using the “ComplexHeatmap’’ package ^87^. For each group and each time period (2017, 2018 separately and 2017-2018 together), GAM model predictions were used to show the direction of responses of richness, evenness, and PD to environmental variables if their relationship was significant in at least one time period (using the “mgcv” package of R ^95^).

Additional multiple linear regression models with the same selection method were run for the changes in local richness (subtracting 2018 values from 2017 values per habitat) and changes in each environmental variable among the two years to link differences in local richness to potential environmental drivers.

All the statistical analysis and illustrations were made with the R program ^96^.

## Acknowledgments

Samples were collected as part of the Interreg V-A Austria-Hungary programme of the European Regional Development Fund (Vogelwarte Madárvárta 2). The authors thank the colleagues of Lake Neusiedl Biological Station, Thomas Zechmeister, Richard Haider, and Rudolf Schalli for their help in field work and for providing access to their laboratories, Christian Preiler, Lilian Lee Müller-Fischer, and Radka Ptacnikova for laboratory analyses and practical help, and Robert Ptacnik for useful discussions and his help during the original project. This work was supported by the RRF-2.3.1-21-2022-00014 project. Beáta Szabó acknowledges support by NKFIH-132095. Zsófia Horváth acknowledges support from the Janos Bolyai Research Scholarship of the Hungarian Academy of Sciences (Grant number BO/00392/20/8) and the Bolyai+ Grant (ÚNKP-22-5-ELTE-84).

## Author contributions

ZH and CFV proposed the original idea; ZH, CFV, ZM collected the samples; ZM performed the laboratory work; ZM, BSz, KP analysed the data; ZM visualized the data; ZM wrote the manuscript with the supervision of ZH; all authors commented on earlier version of the manuscript.

## Additional Information

Competing Interests

The authors declare no competing interests.

## Data Availability

Data analysed during the current study are available in the NCBI SRA repository as part of the BioProject accession PRJNA748202.

## References

1. Céréghino, R., Biggs, J., Oertli, B. & Declerck, S. The ecology of European ponds: defining the characteristics of a neglected freshwater habitat. in Pond Conservation in Europe (eds. Oertli, B. et al.) 1–6 (Springer Netherlands, 2007). doi:10.1007/978-90-481-9088-1_1.

2. Olmo, C. et al. The environmental framework of temporary ponds: A tropical-mediterranean comparison. CATENA 210, 105845 (2022).

3. Griffiths, R. A. Temporary ponds as amphibian habitats. Aquat. Conserv. Mar. Freshw. Ecosyst. 7, 119–126 (1997).

4. Boix, D. et al. Conservation of Temporary Wetlands. in Encyclopedia of the World’s Biomes 279–294 (Elsevier, 2020). doi:10.1016/B978-0-12-409548-9.12003-2.

5. Fritz, K. A. & Whiles, M. R. Reciprocal subsidies between temporary ponds and riparian forests. Limnol. Oceanogr. 66, 3149–3161 (2021).

6. Jeffries, M. The spatial and temporal heterogeneity of macrophyte communities in thirty small, temporary ponds over a period of ten years. Ecography 31, 765–775 (2008).

7. Hassall, C. The ecology and biodiversity of urban ponds. WIREs Water 1, 187–206 (2014).

8. Lukács, B. A. et al. Macrophyte diversity of lakes in the Pannon Ecoregion (Hungary). Limnologica 53, 74–83 (2015).

9. Florencio, M., Díaz-Paniagua, C., Gómez-Rodríguez, C. & Serrano, L. Biodiversity patterns in a macroinvertebrate community of a temporary pond network. Insect Conserv. Divers. 7, 4–21 (2014).

10. Meland, S., Sun, Z., Sokolova, E., Rauch, S. & Brittain, J. E. A comparative study of macroinvertebrate biodiversity in highway stormwater ponds and natural ponds. Sci. Total Environ. 740, 140029 (2020).

11. Hahn, M. W. The microbial diversity of inland waters. Curr. Opin. Biotechnol. 17, 256–261 (2006).

12. Felföldi, T. Microbial communities of soda lakes and pans in the Carpathian Basin: a review. Biol. Futura 71, 393–404 (2020).

13. Grossart, H., Massana, R., McMahon, K. D. & Walsh, D. A. Linking metagenomics to aquatic microbial ecology and biogeochemical cycles. Limnol. Oceanogr. 65, (2020).

14. Marrone, F., Fontaneto, D. & Naselli-Flores, L. Cryptic diversity, niche displacement and our poor understanding of taxonomy and ecology of aquatic microorganisms. Hydrobiologia (2022) doi:10.1007/s10750-022-04904-x.

15. Ducklow, H. jMicrobial services: challenges for microbial ecologists in a changing world. Aquat. Microb. Ecol. 53, 13–19 (2008).

16. Bodelier, P. L. E. Toward understanding, managing, and protecting microbial ecosystems. Front. Microbiol. 2, (2011).

17. Bell, T., Newman, J. A., Silverman, B. W., Turner, S. L. & Lilley, A. K. The contribution of species richness and composition to bacterial services. Nature 436, 1157–1160 (2005).

18. Trivedi, C. et al. Losses in microbial functional diversity reduce the rate of key soil processes. Soil Biol. Biochem. 135, 267–274 (2019).

19. Wellborn, G. A., Skelly, D. K. & Werner, E. E. Mechanisms creating community structure across freshwater habitat gradient. Annu. Rev. Ecol. Syst. 27, 337–363 (1996).

20. Chase, J. M. Drought mediates the importance of stochastic community assembly. Proc. Natl. Acad. Sci. 104, 17430–17434 (2007).

21. Tweed, S., Grace, M., Leblanc, M., Cartwright, I. & Smithyman, D. The individual response of saline lakes to a severe drought. Sci. Total Environ. 409, 3919–3933 (2011).

22. Aguilar, P., Acosta, E., Dorador, C. & Sommaruga, R. Large Differences in bacterial community composition among three nearby extreme waterbodies of the High Andean Plateau. Front. Microbiol. 7, (2016).

23. Boros, E. V. -Balogh, K., Vörös, L. & Horváth, Z. Multiple extreme environmental conditions of intermittent soda pans in the Carpathian Basin (Central Europe). Limnologica 62, 38–46 (2017).

24. Lengyel, E., Pálmai, T., Padisák, J. & Stenger-Kovács, C. Annual hydrological cycle of environmental variables in astatic soda pans (Hungary). J. Hydrol. 575, 1188–1199 (2019).

25. Vieira-Silva, S. & Rocha, E. P. C. The Systemic imprint of growth and its uses in ecological (Meta)genomics. PLoS Genet. 6, e1000808 (2010).

26. Cunillera-Montcusí, D. et al. Freshwater salinisation: a research agenda for a saltier world. Trends Ecol. Evol. 37, 440–453 (2022).

27. Šolić, M. et al. Structure of microbial communities in phosphorus-limited estuaries along the eastern Adriatic coast. J. Mar. Biol. Assoc. U. K. 95, 1565–1578 (2015).

28. Traving, S. J. et al. The Effect of increased loads of dissolved organic matter on estuarine microbial community composition and function. Front. Microbiol. 8, (2017).

29. Zhang, G. et al. Salinity controls soil microbial community structure and function in coastal estuarine wetlands. Environ. Microbiol. 23, 1020–1037 (2021).

30. Tkavc, R. et al. Bacterial communities in the ‘petola’ microbial mat from the Secovlje salterns (Slovenia): Bacterial communities in the ‘petola’. FEMS Microbiol. Ecol. 75, 48–62 (2011).

31. Ali, I. et al. Comparative Study of Physical Factors and Microbial Diversity of Four Man-Made Extreme Ecosystems. Proc. Natl. Acad. Sci. India Sect. B Biol. Sci. 86, 767–778 (2016).

32. Paul, V., Banerjee, Y., Ghosh, P. & Busi, S. B. Depthwise microbiome and isotopic profiling of a moderately saline microbial mat in a solar saltern. Sci. Rep. 10, 20686 (2020).

33. Stenger-Kovács, C. et al. Vanishing world: alkaline, saline lakes in Central Europe and their diatom assemblages. Inland Waters 4, 383–396 (2014).

34. Stenger-Kovács, C., Hajnal, É., Lengyel, E., Buczkó, K. & Padisák, J. A test of traditional diversity measures and taxonomic distinctness indices on benthic diatoms of soda pans in the Carpathian basin. Ecol. Indic. 64, 1–8 (2016).

35. Szabó, B., Lengyel, E., Padisák, J., Vass, M. & Stenger-Kovács, C. Structuring forces and β-diversity of benthic diatom metacommunities in soda pans of the Carpathian Basin. Eur. J. Phycol. 53, 219–229 (2018).

36. Szabó, A. et al. Soda pans of the Pannonian steppe harbor unique bacterial communities adapted to multiple extreme conditions. Extremophiles 21, 639–649 (2017).

37. Szabó, A. et al. Grazing pressure-induced shift in planktonic bacterial communities with the dominance of acIII-A1 actinobacterial lineage in soda pans. Sci. Rep. 10, 19871 (2020).

38. Benlloch, S. et al. Prokaryotic genetic diversity throughout the salinity gradient of a coastal solar saltern. Environ. Microbiol. 4, 349–360 (2002).

39. Horváth, Z. et al. Opposing patterns of zooplankton diversity and functioning along a natural stress gradient: when the going gets tough, the tough get going. Oikos 123, 461–471 (2014).

40. Mo, Y. et al. Low shifts in salinity determined assembly processes and network stability of microeukaryotic plankton communities in a subtropical urban reservoir. Microbiome 9, 128 (2021).

41. Gómez-Rodríguez, C., Bustamante, J. & Díaz-Paniagua, C. Evidence of hydroperiod shortening in a preserved system of temporary ponds. Remote Sens. 2, 1439–1462 (2010).

42. Finger Higgens, R. A. et al. Changing lake dynamics indicate a drier arctic in western greenland. J. Geophys. Res. Biogeosciences 124, 870–883 (2019).

43. Pekel, J.-F., Cottam, A., Gorelick, N. & Belward, A. S. High-resolution mapping of global surface water and its long-term changes. Nature 540, 418–422 (2016).

44. Grillas, P., Rhazi, L., Lefebvre, G., El Madihi, M. & Poulin, B. Foreseen impact of climate change on temporary ponds located along a latitudinal gradient in Morocco. Inland Waters 11, 492–507 (2021).

45. Xi, Y., Peng, S., Ciais, P. & Chen, Y. Future impacts of climate change on inland Ramsar wetlands. Nat. Clim. Change 11, 45–51 (2021).

46. Zacharias, I. & Zamparas, M. Mediterranean temporary ponds. A disappearing ecosystem. Biodivers. Conserv. 19, 3827–3834 (2010).

47. Horváth, Z., Ptacnik, R., Vad, C. F. & Chase, J. M. Habitat loss over six decades accelerates regional and local biodiversity loss via changing landscape connectance. Ecol. Lett. 22, 1019–1027 (2019).

48. Zhong, Y. et al. Shrinking habitats and native species loss under climate change: a multifactorial risk Assessment of china’s inland wetlands. 28 (2022).

49. Whiting, G. J. & Chanton, J. P. Greenhouse carbon balance of wetlands: methane emission versus carbon sequestration: Greenhouse carbon balance of wetlands. Tellus B 53, 521–528 (2001).

50. Mitsch, W. J. et al. Wetlands, carbon, and climate change. Landsc. Ecol. 28, 583–597 (2013).

51. Ardón, M., Helton, A. M. & Bernhardt, E. S. Salinity effects on greenhouse gas emissions from wetland soils are contingent upon hydrologic setting: a microcosm experiment. Biogeochemistry 140, 217–232 (2018).

52. Jeppesen, E., Beklioğlu, M., Özkan, K. & Akyürek, Z. Salinization increase due to climate change will have substantial negative effects on inland waters: A call for multifaceted research at the local and global scale. The Innovation 1, 100030 (2020).

53. Boros, E., Horváth, Zs., Wolfram, G. & Vörös, L. Salinity and ionic composition of the shallow astatic soda pans in the Carpathian Basin. Ann. Limnol. - Int. J. Limnol. 50, 59–69 (2014).

54. Williams, D. D. The Ecology of Temporary Waters. (Springer Netherlands, 1987). doi:10.1007/978-94-011-6084-1.

55. Hammer, U. T. The effects of climate change on the salinity, water levels and biota of Canadian prairie saline lakes. SIL Proc. 1922-2010 24, 321–326 (1990).

56. Schallenberg, M., Hall, C. & Burns, C. Consequences of climate-induced salinity increases on zooplankton abundance and diversity in coastal lakes. Mar. Ecol. Prog. Ser. 251, 181–189 (2003).

57. Felföldi, T., Somogyi, B., Márialigeti, K. & Vörös, L. Characterization of photoautotrophic picoplankton assemblages in turbid, alkaline lakes of the Carpathian Basin (Central Europe). J. Limnol. 68, 385 (2009).

58. Somogyi, B. et al. Winter bloom of picoeukaryotes in Hungarian shallow turbid soda pans and the role of light and temperature. Aquat. Ecol. 43, 735–744 (2009).

59. Pálffy, K. et al. Unique picoeukaryotic algal community under multiple environmental stress conditions in a shallow, alkaline pan. Extremophiles 18, 111–119 (2014).

60. Padisák, J. & Naselli-Flores, L. Phytoplankton in extreme environments: importance and consequences of habitat permanency. Hydrobiologia 848, 157–176 (2021).

61. Olli, K., Ptacnik, R., Klais, R. & Tamminen, T. Phytoplankton species richness along coastal and estuarine salinity continua. Am. Nat. 194, E41–E51 (2019).

62. Olli, K., Tamminen, T. & Ptacnik, R. Predictable shifts in diversity and ecosystem function in phytoplankton communities along coastal salinity continua. Limnol. Oceanogr. Lett. lol2.10242 (2022) doi:10.1002/lol2.10242.

63. Tikhonenkov, D. V., Burkovsky, I. V. & Mazei, Y. A. Is there a relation between the distribution of heterotrophic flagellates and the zonation of a marine intertidal flat? 55, 13 (2015).

64. Hartmut Arndt et al. Functional diversity of heterotrophic flagellates in aquatic ecosystems. in Flagellates 252–280 (CRC Press, 2000). doi:10.1201/9781482268225-18.

65. Je Lee, W. & Patterson, D. J. Diversity and geographic distribution of free-living heterotrophic flagellates – Analysis by PRIMER. Protist 149, 229–244 (1998).

66. Azovsky, A. I., Tikhonenkov, D. V. & Mazei, Y. A. An estimation of the global diversity and distribution of the smallest eukaryotes: biogeography of marine benthic heterotrophic flagellates. Protist 167, 411–424 (2016).

67. Tikhonenkov, D. V., Mazei, Y. A. & Mylnikov, A. P. Species diversity of heterotrophic flagellates in White Sea littoral sites. Eur. J. Protistol. 42, 191–200 (2006).

68. Van der Gucht, K. et al. The power of species sorting: Local factors drive bacterial community composition over a wide range of spatial scales. Proc. Natl. Acad. Sci. 104, 20404–20409 (2007).

69. Vanschoenwinkel, B. et al. Species sorting in space and time—the impact of disturbance regime on community assembly in a temporary pool metacommunity. J. North Am. Benthol. Soc. 29, 1267–1278 (2010).

70. Datry, T. et al. Metacommunity patterns across three neotropical catchments with varying environmental harshness. Freshw. Biol. 61, 277–292 (2016).

71. Horváth, Z., Vad, C. F., Vörös, L. & Boros, E. The keystone role of anostracans and copepods in European soda pans during the spring migration of waterbirds: The keystone trophic role of crustaceans in European soda pans. Freshw. Biol. 58, 430–440 (2013).

72. Stenger-Kovács, C. & Lengyel, E. Taxonomical and distribution guide of diatoms in soda pans of Central Europe. Stud. Bot. Hung. 46, 3–203 (2015).

73. Szabó, B. et al. Microbial stowaways: Waterbirds as dispersal vectors of aquatic pro-and microeukaryotic communities. J. Biogeogr. 49, 1286–1298 (2022).

74. Sorokin, D. Y. et al. Microbial diversity and biogeochemical cycling in soda lakes. Extremophiles 18, 791–809 (2014).

75. Hansen, H. P. & Koroleff, F. Determination of nutrients. in Methods of Seawater Analysis (eds. Grasshoff, K., Kremling, K. & Ehrhardt, M.) 159–228 (Wiley-VCH Verlag GmbH, 1999). doi:10.1002/9783527613984.ch10.

76. Parada, A. E., Needham, D. M. & Fuhrman, J. A. Every base matters: assessing small subunit rRNA primers for marine microbiomes with mock communities, time series and global field samples: Primers for marine microbiome studies. Environ. Microbiol. 18, 1403–1414 (2016).

77. Apprill, A., McNally, S., Parsons, R. & Weber, L. Minor revision to V4 region SSU rRNA 806R gene primer greatly increases detection of SAR11 bacterioplankton. Aquat. Microb. Ecol. 75, 129–137 (2015).

78. Ray, J. L. et al. Metabarcoding and metabolome analyses of copepod grazing reveal feeding preference and linkage to metabolite classes in dynamic microbial plankton communities. Mol. Ecol. 25, 5585–5602 (2016).

79. Schloss, P. D. et al. Introducing mothur: Open-Source, Platform-Independent, Community-Supported Software for Describing and Comparing Microbial Communities. Appl. Environ. Microbiol. 75, 7537–7541 (2009).

80. Kozich, J. J., Westcott, S. L., Baxter, N. T., Highlander, S. K. & Schloss, P. D. Development of a dual-index sequencing strategy and curation pipeline for analyzing amplicon sequence sata on the MiSeq Illumina sequencing platform. Appl. Environ. Microbiol. 79, 5112–5120 (2013).

81. Kunin, V., Engelbrektson, A., Ochman, H. & Hugenholtz, P. Wrinkles in the rare biosphere: pyrosequencing errors can lead to artificial inflation of diversity estimates. Environ. Microbiol. 12, 118–123 (2010).

82. Quast, C. et al. The SILVA ribosomal RNA gene database project: improved data processing and web-based tools. Nucleic Acids Res. 41, D590–D596 (2012).

83. Rohwer, R. R., Hamilton, J. J., Newton, R. J. & McMahon, K. D. TaxAss: Leveraging a Custom Freshwater Database Achieves Fine-Scale Taxonomic Resolution. mSphere 3, e00327–18 (2018).

84. Guillou, L. et al. The Protist Ribosomal Reference database (PR2): a catalog of unicellular eukaryote Small Sub-Unit rRNA sequences with curated taxonomy. Nucleic Acids Res. 41, D597–D604 (2012).

85. McMurdie, P. J. & Holmes, S. phyloseq: An R package for reproducible interactive analysis and graphics of microbiome census data. PLoS ONE 8, e61217 (2013).

86. Burian, A. et al. Predation increases multiple components of microbial diversity in activated sludge communities. ISME J. 16, 1086–1094 (2022).

87. Gu, Z. Complex heatmap visualization. iMeta 1, (2022).

88. Atkinson, S. T. et al. Substantial long-term loss of alpha and gamma diversity of lake invertebrates in a landscape exposed to a drying climate. Glob. Change Biol. 27, 6263–6279 (2021).

89. Clesceri LS, Greenberg AE, Eaton AD. 1999. Standard methods for examination of water and wastewater (20th ed). Available from: http://ipkosar.ir/jspui/handle/961944/280820

90. vegan: Community Ecology Package. R package version 2.5–7.: Jari Oksanen, F. Guillaume Blanchet, Michael Friendly, Roeland Kindt, Pierre Legendre, Dan McGlinn, Peter, R. Minchin, R. B. O’Hara Gavin L. Simpson Peter Solymos, M. Henry H. Stevens Eduard Szoecs and Helene Wagner (2020). https://CRAN.R-project.org/package=vegan

91. ggpubr R package version 0.4.0.: Alboukadel Kassambara (2020). ggpubr: ‘ggplot2’ Based Publication Ready Plots. https://CRAN.R-project.org/package=ggpubr

92. pairwiseAdonis R package version 0.4.: Pedro Martinez Arbizu (2017). pairwiseAdonis: Pairwise Multilevel Comparison using Adonis. R package. https://cran.r-project.org/web/packages/pairwise/index.html

93. picante R package version 1.8.2.: S.W. Kembel, P.D. Cowan, M.R. Helmus, W.K. Cornwell, H. Morlon, D.D. Ackerly, S.P. Blomberg, and C.O. Webb. 2010. Picante: R tools for integrating phylogenies and ecology. Bioinformatics 26:1463–1464. https://cran.r-project.org/web/packages/picante/index.html

94. MASS R package version 7.3-54. Venables, W. N. & Ripley, B. D. (2002) Modern Applied Statistics with S. Fourth Edition. Springer, New York. ISBN 0-387-95457-0. https://cran.r-project.org/web/packages/MASS/index.html

95. mgcv R package version 1.8-38.: Wood, S.N. (2011) Fast stable restricted maximum likelihood and marginal likelihood estimation of semiparametric generalized linear models. Journal of the Royal Statistical Society (B) 73(1):3–36. https://cran.r-project.org/web/packages/mgcv/index.html

96. R Core Team (2022). R: A language and environment for statistical computing. R Foundation for Statistical Computing, Vienna, Austria. https://www.R-project.org/.

97. Hadziavdic, K. et al. Characterization of the 18S rRNA Gene for Designing Universal Eukaryote Specific Primers. PLoS ONE. 9, e87624 (2014).

98. Horváth, Z., Vad, C. F., Vörös, L. & Boros, E. The keystone role of anostracans and copepods in European soda pans during the spring migration of waterbirds: The keystone trophic role of crustaceans in European soda pans. Freshw. Biol. 58, 430–440 (2013).

