## Supplementary material for "Environmental changes associated with drying climate are expected to affect functional groups of pro- and microeukaryotes differently in temporary saline waters"


This file includes the following text, tables and figures:

Text S1

Table S1-S6

Figures S1-S9

Text S1 Description of the generalized additive models of the meteorological data of the Seewinkel region

Generalized additive models (GAM) with smooth terms (using the “mgcv” package of R, Wood, 2017) were used to analyze the temporal trends of the annual mean temperature and annual precipitation data from 2009 (downloaded: 2022.01.12., ZAMG - Zentralanstalt für Meteorologie und Geodynamik, Station Eisenstadt ) till 2020. Six models were created for each meteorological data with different numbers of knots (2-7), and model selection was based on the Akaike’s Information Criteria (AIC). An additional final model check was done with the gam_check function of the “mgcv” package of R. For the annual mean temperature, the model with 2 knots and for the annual mean precipitation, the model with 3 knots were the best fitted models.

Reference of Text S1

Wood, S. N. (2011) Fast stable restricted maximum likelihood and marginal likelihood estimation of semiparametric generalized linear models. Journal of the Royal Statistical Society (B) 73 (1):3-36

Table S1 GPS coordinates of the sampling sites in the Seewinkel region and the sampling years we collected samples from each site (black dots)

| Name | Longitude | Latitude | 2017 | 2018 |
|---|---|---|---|---|
| Albersee | 16.77011917 | 47.77513639 | ● | ● |
| Apetloner Meierhoflacke | 16.82404028 | 47.72157583 | | ● |
| Auerlacke | 16.88677778 | 47.78917083 | ● | ● |
| Birnbaumlacke | 16.86491056 | 47.81771528 | ● | ● |
| Grosse Neubruchlacke | 16.84219833 | 47.78612111 | ● | ● |
| Herrnsee | 16.7699880 | 47.7445640 | | ● |
| Kirchsee | 16.78548861 | 47.75867861 | | ● |
| Krautingsee | 16.78047972 | 47.75602861 | ● | ● |
| Kühnbrunnlacke | 16.87862583 | 47.79273556 | ● | ● |
| Lange Lacke | 16.87876472 | 47.75747528 | ● | ● |
| Martenhofenlacke | 16.8566950 | 47.75050167 | ● | ● |
| Mittlerer Stinkersee | 16.78752194 | 47.80675889 | ● | ● |
| Neufeldlacke | 16.84024833 | 47.76447667 | ● | ● |
| Unnamed | 16.8443250 | 47.7680360 | ● | |
| Obere Höllacke | 16.80793222 | 47.82711944 | ● | ● |
| Oberer Stinkersee | 16.79251722 | 47.81376222 | ● | ● |
| Ochsebrunnlacke | 16.84461722 | 47.81073583 | ● | ● |
| Östliche Wörthenlacke | 16.88009389 | 47.77334361 | ● | ● |
| Östliche Fuchslochlacke | 16.86618583 | 47.79044083 | ● | ● |
| Runde Lacke | 16.79274361 | 47.78574472 | ● | ● |
| Sechsmahdlacke | 16.88411194 | 47.78378861 | ● | ● |
| Stundlacke | 16.87170639 | 47.79890389 | ● | ● |
| Südlicher Silbersee | 16.77973694 | 47.79113278 | ● | ● |
| Westliche Wörthenlacke | 16.87077667 | 47.77092167 | ● | ● |
| Westliche Fuschslochlacke | 16.85234944 | 47.79007667 | ● | ● |
| Zicklacke | 16.78475194 | 47.76696444 | ● | ● |

Table S2 Meteorological data of the Seewinkel region four months prior to the sampling in April 2017 and April 2018 (source: ZAMG - Zentralanstalt für Meteorologie und Geodynamik, Station Eisenstadt)

| Year | Month | Average temperature (°C) | Amount of precipitation (mm) | Sunshine duration (hour) | Average precipitation in the four months (mm) |
|---|---|---|---|---|---|
| 2016 | December | 1.20 | 20 | 90 | 26 |
| 2017 | January | -3.7 | 14 | 95 | |
| 2017 | February | 3 | 44 | 83 | |
| 2017 | March | 9.20 | 34 | 196 | |
| 2017 | December | 3 | 57 | 82 | 47 |
| 2018 | January | 3.80 | 33 | 58 | |
| 2018 | February | -0.8 | 45 | 75 | |
| 2018 | March | 3.20 | 53 | 117 | |

Table S3 Amplification of the 16S rRNA and 18 rRNA genes was carried out by LGC Genomics (Berlin, Germany). The pooled PCR products were purified with Agencourt AMPure XP beads (Beckman Coulter, Inc., IN, USA) to remove primer dimer, and an additional purification on MiniElute columns (QIAGEN GmbH, Hilden, Germany) was carried out. About 100 ng of each DNA pool was used to prepare the Illumina libraries using the Ovation Rapid DR Multiplex System 1-96 (NuGEN Technologies, Inc., CA, USA). Illumina libraries were pooled (Illumina, Inc., CA, USA) and size was selected by preparative gel electrophoresis. Sequencing was performed on an Illumina MiSeq platform

| **Polymerase chain reaction mastermix** | **1x** |
|---|---|
| 1 x MyTaq buffer | 20 µL |
| MyTaq DNA polymerase (Bioline GmbH, Luckenwalde, Germany) | 1.5 units |
| BioStabII PCR Enhancer (Sigma-Aldrich Co.) | 2 µl |
| Forward primer | 15 pmol |
| Reverse primer | 15 pmol |
| DNA (1-10 ng) | 1 µl |

| **Heat cycle of the PCR (30 cycles)** | **Temperature (°C)** | **Duration** |
|---|---|---|
| Pre-denaturation | 96°C | 1 min |
| Denaturation | 96°C | 15 s |
| Annealing | 55 °C | 30 s |
| Extension | 70 °C | 90 s |
| Hold | 8 °C | ∞ |

Table S4 Interset correlations (Pearson's correlation coefficients) between the environmental variables and the canonical axes of the CCA of prokaryotic communities (bold: significant environmental variable for the community composition)

| **Environmental variables** | **CCA1** | **CCA2** |
|---|---|---|
| **2017** | | |
| **Water depth** | **-0.52** | **0.57** |
| pH | 0.00 | 0.08 |
| Conductivity | -0.19 | 0.19 |
| **TP** | **0.96** | **0.04** |
| TN | 0.51 | -0.58 |
| TSS | 0.93 | 0.24 |
| **2018** | | |
| **Water depth** | **-0.52** | **0.51** |
| pH | -0.50 | 0.57 |
| **Conductivity** | **-0.66** | **0.59** |
| **TP** | **0.75** | **0.30** |
| TN | 0.17 | 0.59 |
| TSS | 0.79 | 0.03 |

Table S5 Interset correlations (Pearson's correlation coefficients) between the environmental variables and the canonical axes of the CCA of microeukaryotic communities (bold: significant environmental variable for the community composition)

| **Environmental variables** | **CCA1** | **CCA2** |
|---|---|---|
| **2017** | | |
| Water depth | 0.09 | -0.31 |
| pH | -0.62 | -0.12 |
| **Conductivity** | **-0.70** | **-0.54** |
| **TP** | **-0.58** | **0.72** |
| TN | -0.28 | 0.52 |
| TSS | -0.60 | 0.70 |
| **2018** | | |
| Water depth | 0.22 | 0.35 |
| **pH** | **0.88** | **0.15** |
| **Conductivity** | **0.80** | **-0.10** |
| TP | 0.27 | -0.36 |
| TN | 0.55 | -0.19 |
| TSS | 0.15 | -0.06 |

Table S6 Summary statistics of the multiple linear regression models testing the relationships between richness change and environmental variables change from 2017 to 2018

(*= p < 0.01 , NS = not significant)

| Group | AIC | $R^2$ | p (model) | Significance | p | Direction of relationship |
|---|---|---|---|---|---|---|
| **Bacteria** | Conductivity | 0.153 | 0.045 | * | 0.045 | - |
| **Cyanobacteria** | Conductivity | 0.147 | 0.129 | NS | 0.111 | |
| | TP | 0.147 | 0.129 | NS | 0.177 | |
| **Ciliates** | Conductivity | 0.249 | 0.064 | * | 0.027 | - |
| | TP | 0.249 | 0.064 | NS | 0.108 | |
| | TSS | 0.249 | 0.064 | NS | 0.190 | |
| **Fungi** | pH | 0.274 | 0.013 | * | 0.013 | - |
| **HF-HNF** | Water depth | 0.351 | 0.265 | NS | 0.104 | |
| | pH | 0.351 | 0.265 | NS | 0.382 | |
| | TP | 0.351 | 0.265 | NS | 0.319 | |
| | TN | 0.351 | 0.265 | NS | 0.105 | |
| | TSS | 0.351 | 0.265 | NS | 0.119 | |
| **Phytoplankton** | Water depth | -0.349 | 0.962 | NS | 0.493 | |
| | Conductivity | -0.349 | 0.962 | NS | 0.531 | |
| | pH | -0.349 | 0.962 | NS | 0.916 | |
| | TP | -0.349 | 0.962 | NS | 0.996 | |
| | TN | -0.349 | 0.962 | NS | 0.586 | |
| | TSS | -0.349 | 0.962 | NS | 0.802 | |

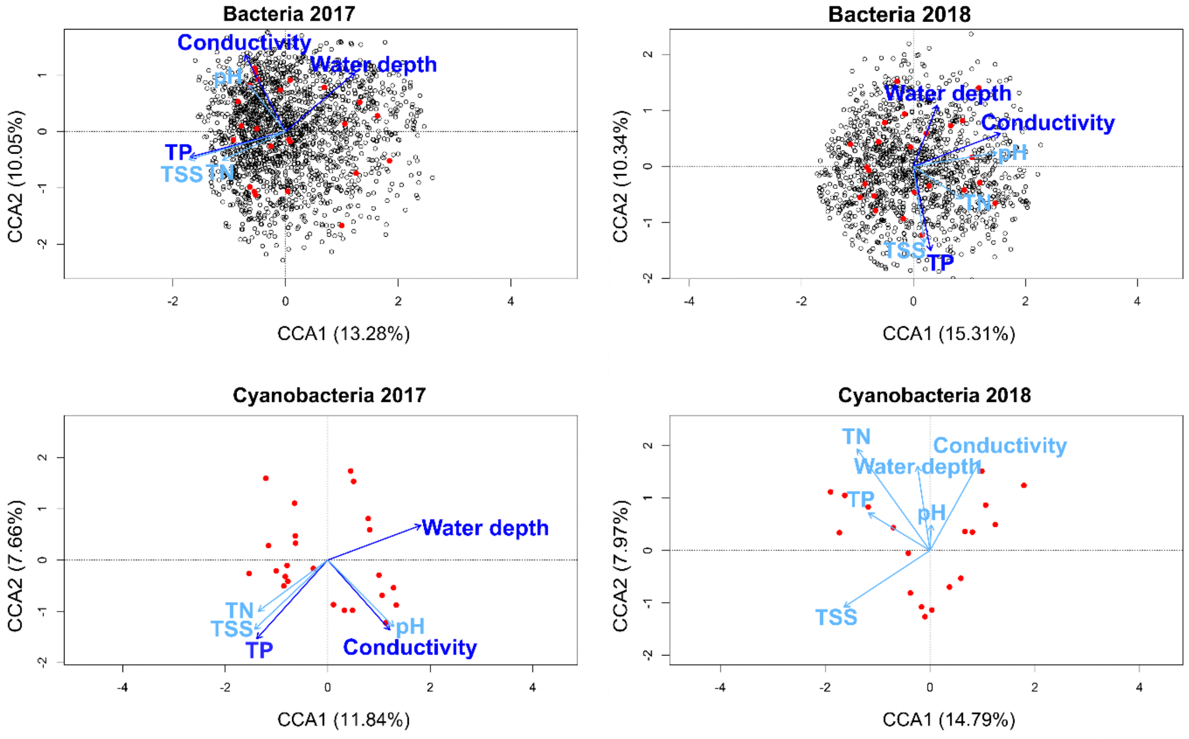


Figure S1 CCAs based on OTU composition of the prokaryotes (red points: habitat scores, dark blue arrows: significant environmental variables, light blue arrows: all other environmental variables)

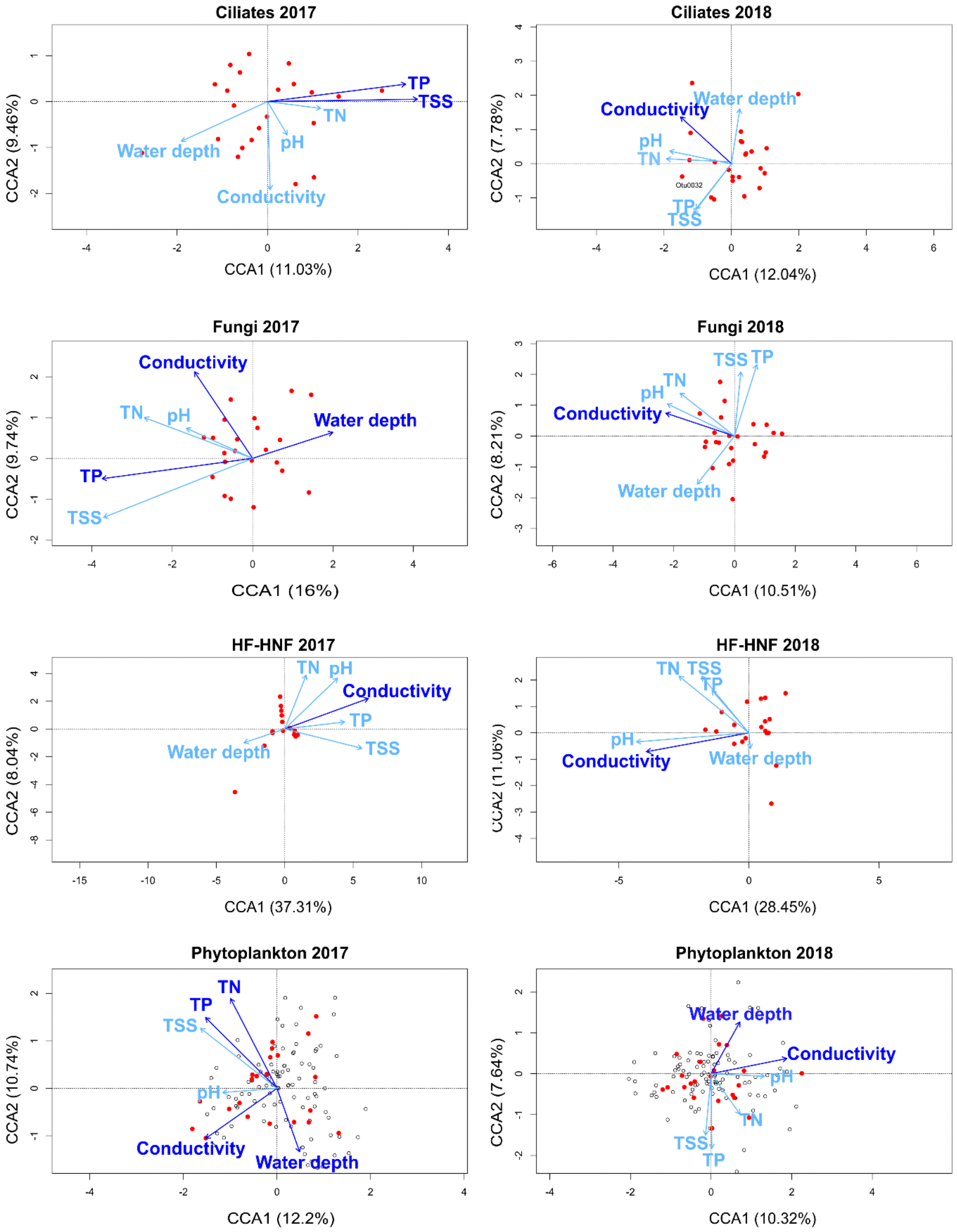


Figure S2 CCAs based on the OTU composition of the microeukaryotes (red points: habitat scores, dark blue arrows: significant environmental variables, light blue arrows: all other environmental variables)

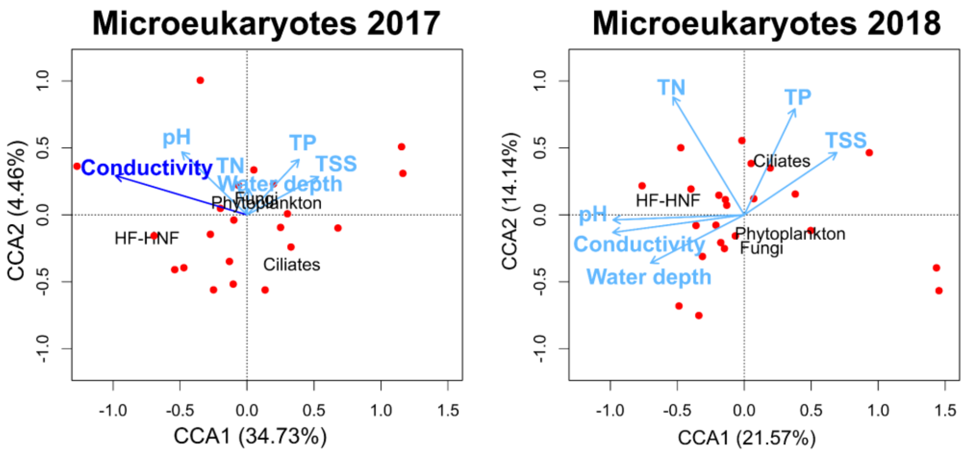


Figure S3 CCA biplots based on the relative abundance of the microeukaryotes (red points: habitat scores, dark blue arrows: significant environmental variables, light blue arrows: all other environmental variables)

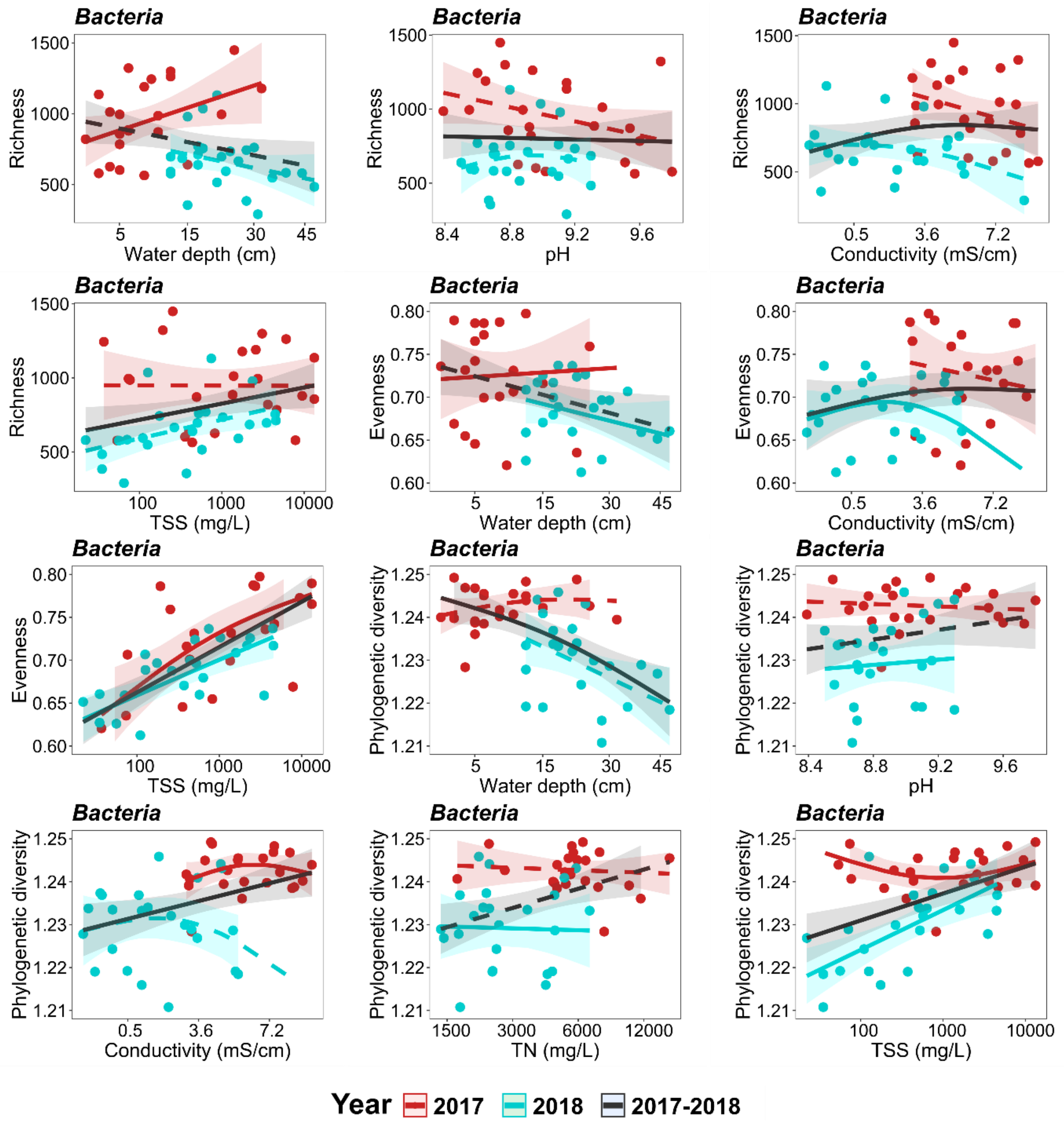


Figure S4 Relationship of richness, evenness, and phylogenetic diversity with the significant environmental variables. Fitted trend lines are based on predicted GAM models in 2017, 2018, and for the pooled dataset. Solid lines: significant relationships, dashed lines: not significant ($p < 0.05$)

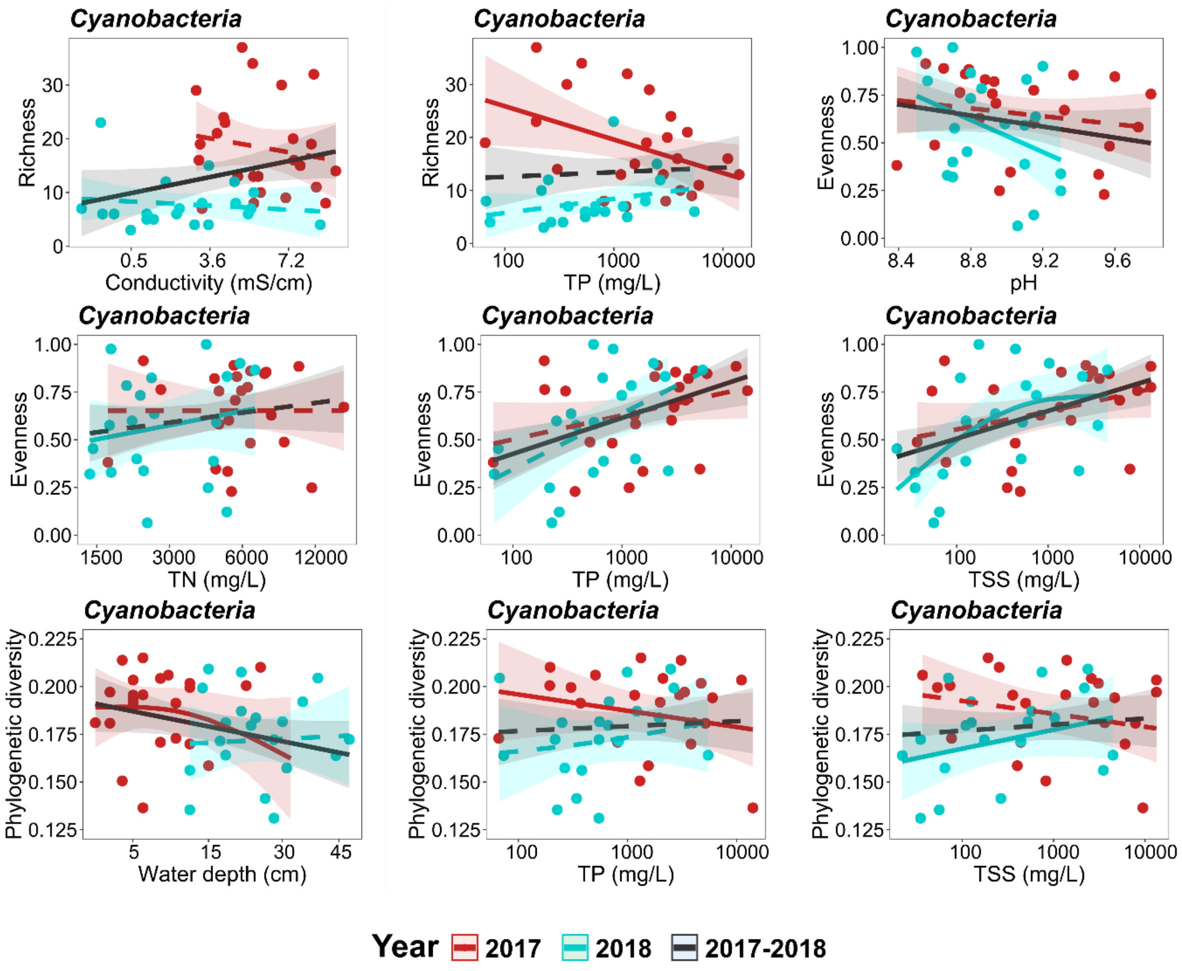


Figure S5 Relationship of richness, evenness, and phylogenetic diversity with the significant environmental variables. Fitted trend lines are based on predicted GAM models in 2017, 2018, and for the pooled dataset. Solid lines: significant relationships, dashed lines: not significant ($p < 0.05$)

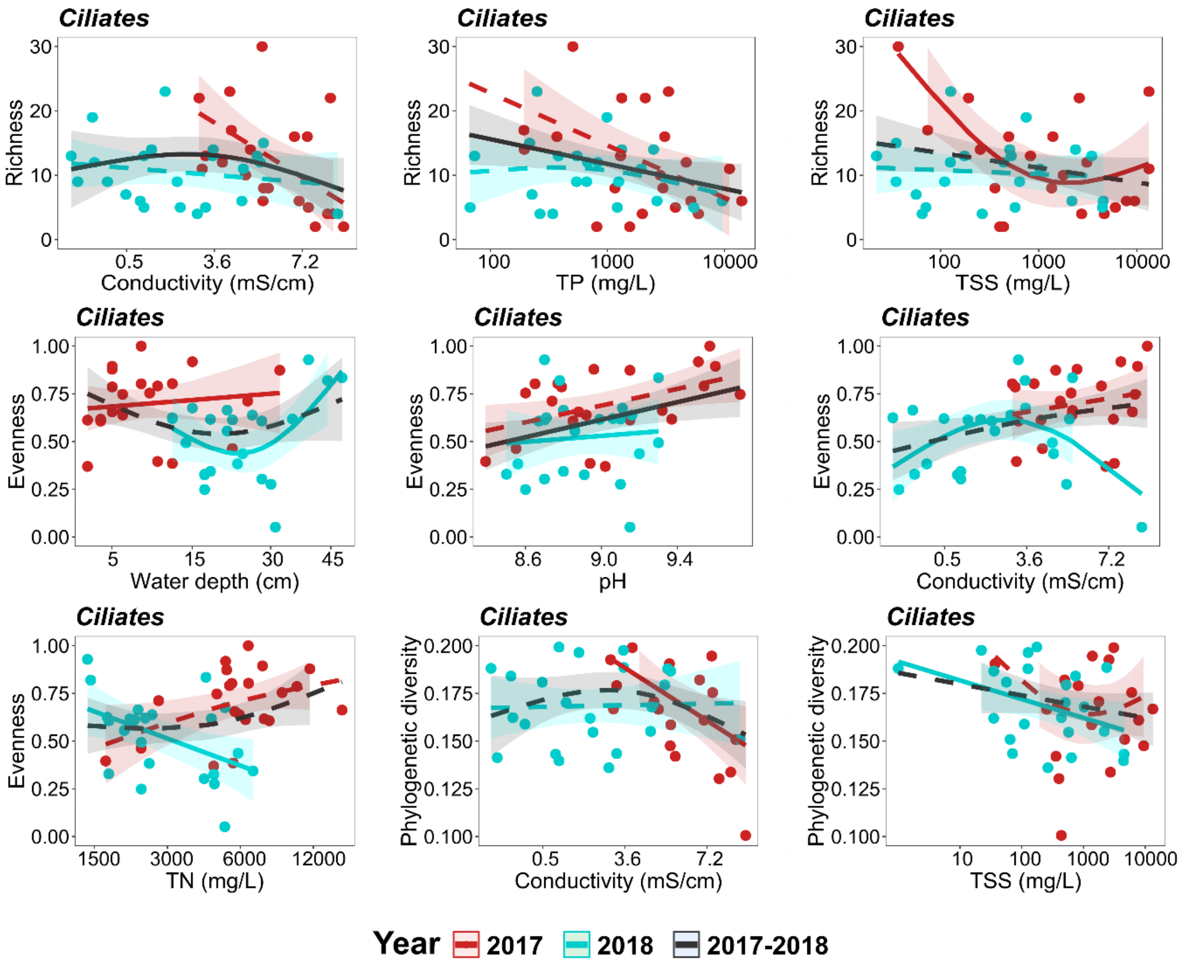


Figure S6 Relationship of richness, evenness, and phylogenetic diversity with the significant environmental variables. Fitted trend lines are based on predicted GAM models in 2017, 2018, and for the pooled dataset. Solid lines: significant relationships, dashed lines: not significant ($p < 0.05$)

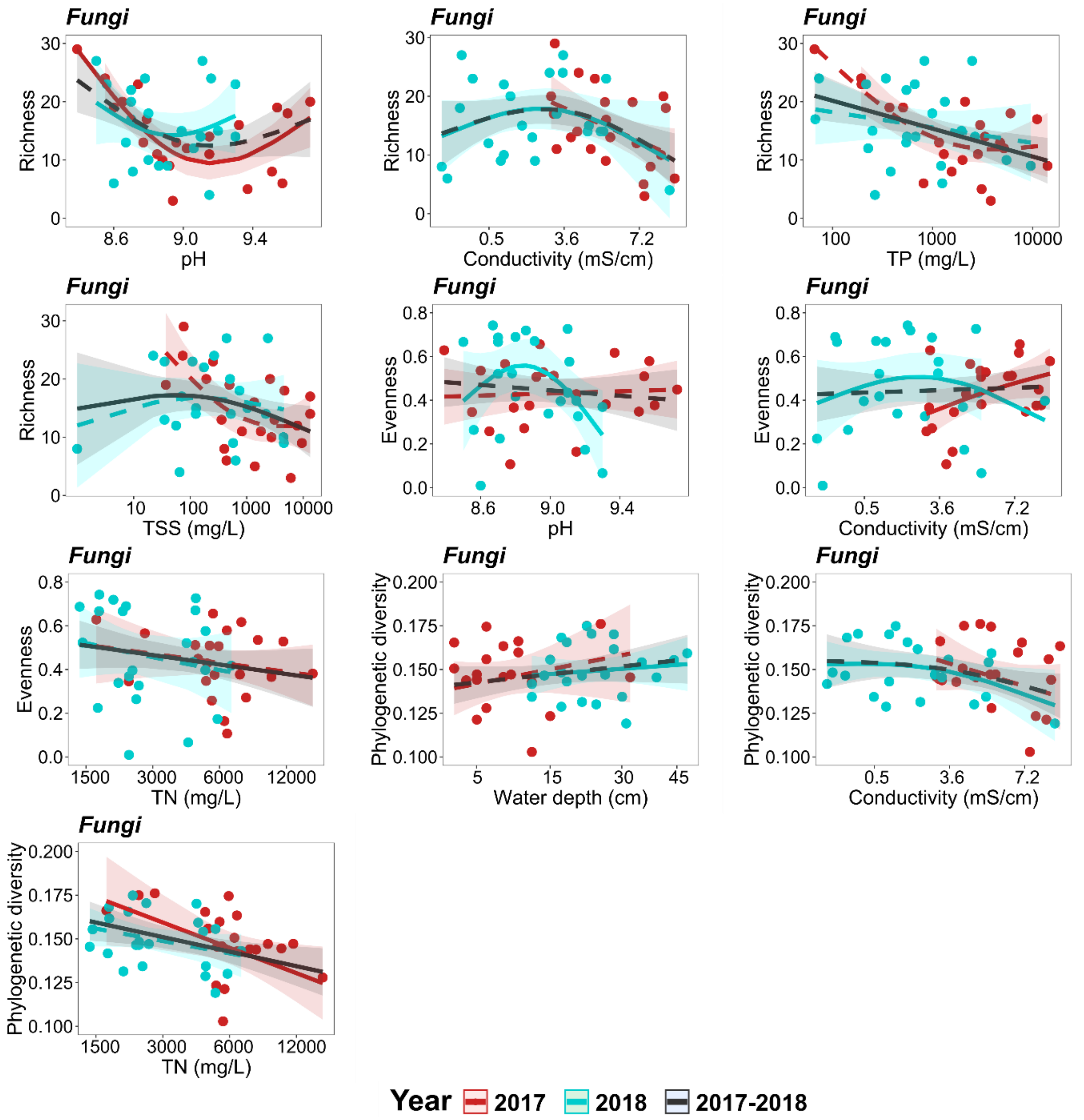


Figure S7 Relationship of richness, evenness, and phylogenetic diversity with the significant environmental variables. Fitted trend lines are based on predicted GAM models in 2017, 2018, and for the pooled dataset. Solid lines: significant relationships, dashed lines: not significant ($p < 0.05$)

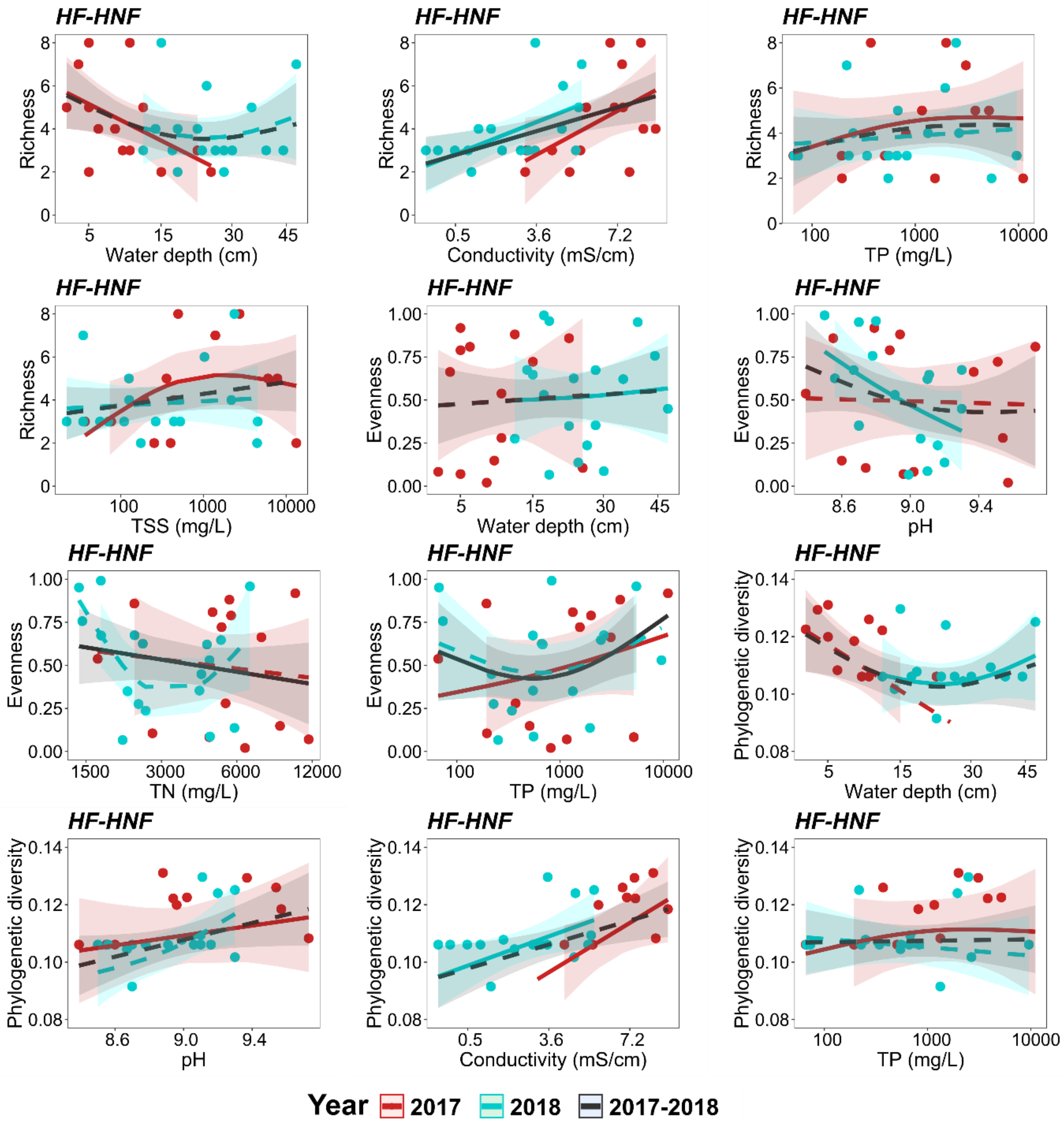


Figure S8 Relationship of richness, evenness, and phylogenetic diversity with the significant environmental variables. Fitted trend lines are based on predicted GAM models in 2017, 2018, and for the pooled dataset. Solid lines: significant relationships, dashed lines: not significant ($p < 0.05$)

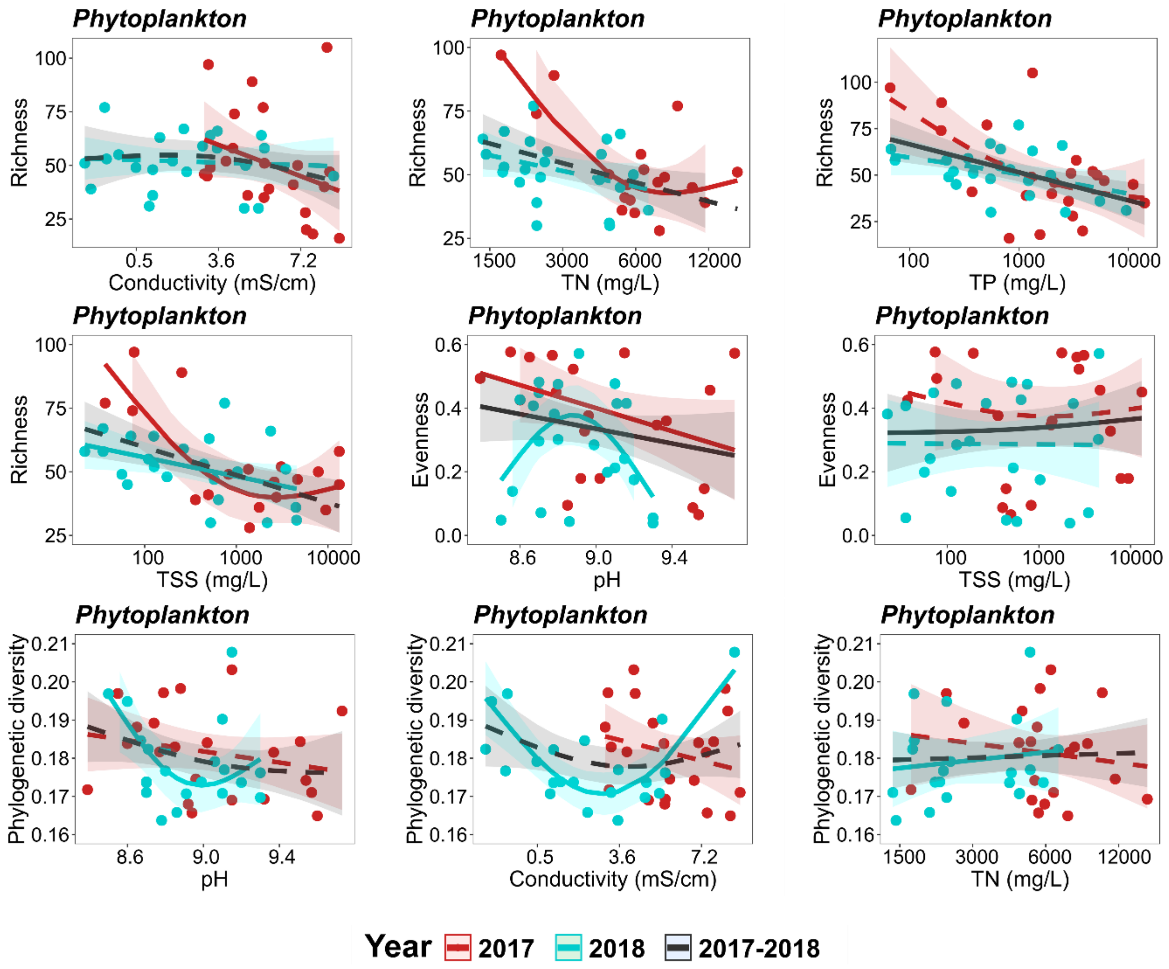


Figure S9 Relationship of richness, evenness, and phylogenetic diversity with the significant environmental variables. Fitted trend lines are based on predicted GAM models in 2017, 2018, and for the pooled dataset. Solid lines: significant relationships, dashed lines: not significant ($p < 0.05$)
